# Determinants of survival and dispersal along the range expansion of a biological invasion

**DOI:** 10.1101/2022.09.14.507789

**Authors:** Eric Edeline, Agnès Starck, Yoann Bennevault, Jean-Marc Paillisson, Eric J. Petit

## Abstract

Projecting and managing the future response of biological systems to global change requires a mechanistic understanding of how climate and ecology jointly drive species demography and range dynamics. Such knowledge is particularly crucial when it comes to invasive species, which expansion may have far-reaching consequences for recipient ecosystems. Here, we use mark recapture in replicated outdoor mesocosms to examine how survival and dispersal, two key drivers of population and range dynamics, respond to climate and ecology in the invasive red swamp crayfish (*Procambarus clarkii*) along an invasion gradient. We show that crayfish survival probability increased with (*i*) increasing body size at high (but not low) crayfish density and (*ii*) with warmer temperatures, and decreased (*i*) with increasing body condition and (*ii*) under higher crayfish density. Overland dispersal probability by crayfish increased with increasing (*i*) body-size, (*ii*) body condition and (*iii*) temperatures. In contrast, crayfish from range-edge and range-core habitats had similar survival and overland dispersal probabilities, suggesting no evolution of the crayfish expansion potential along the invasion gradient. Our results highlight that species population dynamics and range shifts in a changing world are driven by joint contributions from both climate and ecology. In *P. clarkii*, global warming will simultaneously promote both a demographic increase and a geographic range expansion, especially in populations dominated by large-bodied individuals. In already-invaded ecosystems, selective harvesting of large-bodied crayfish can potentially reduce the dispersal potential of populations and, after a few generations, might further induce an evolutionary decline in fitness traits that is desirable from a management perspective.

**Open research statement:** Upon acceptance of this manuscript, data and codes will be made publicly available online on the INRAE data repository (https://entrepot.recherche.data.gouv.fr/dataverse/inrae).

## 1. INTRODUCTION

How species geographic distributions will change in a near future is a central question in ecology, evolution, biodiversity conservation and the management of biological invasions. To date, our ability to answer this question often rests on correlative species distribution models, which provide a poor mechanistic understanding and, arguably, have a low power to predict how organisms will respond to global change. A recent approach to addressing these questions consists in explicitly modelling the ecological and evolutionary processes that control geographic distributions in so called process-explicit models (Briscoe et al., 2019; Townsend Peterson et al., 2015; Travis and Dytham, 2012). These models draw heavily on population-dynamic models, and on how demographic rates, i.e., survival, reproduction and dispersal, respond to environmental change (Fig. 1).

**Fig. 1.**
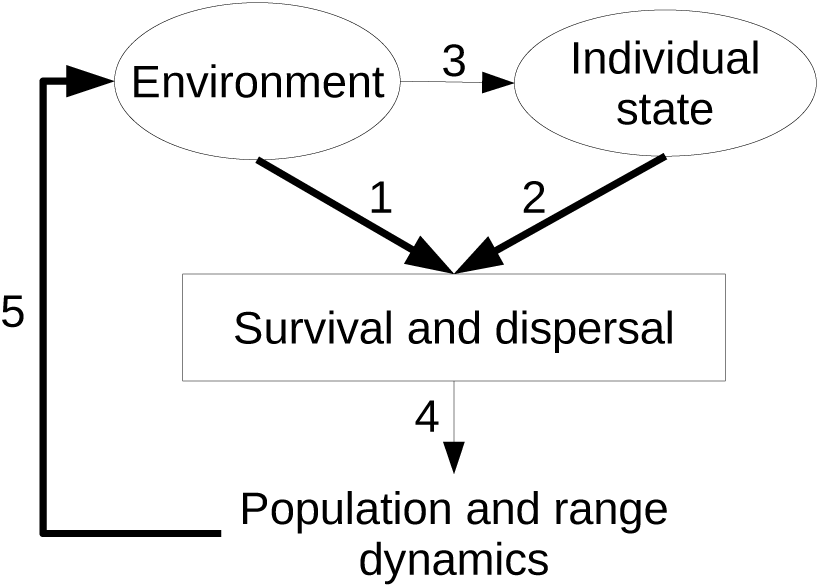
Conceptual framework for process-explicit models of species range dynamics. The environment influences demographic rates either directly (Arrow 1), or indirectly through altering individual state variables through both phenotypic plasticity and the selection experienced by individuals (Arrow 2-3 sequence). Demographic rates ultimately drive the dynamics of both population sizes and geographic ranges (Arrow 4). In turn, population and range dynamics entail environmental alterations (Arrow 5). Bold arrows show the causality links that are specifically investigated in the present study.

In taking a process-explicit approach, the complexity of mechanistically predicting how demographic rates respond to environmental change may be usefully tackled by separating the changes that act directly onto demographic rates, or indirectly through a modification of individual state variables (Fig. 1).

A host of environmental abiotic (e.g., climate) and ecological (i.e., density-dependent) factors have long been known to directly influence demographic rates, including survival and dispersal (Arrow 1 in Fig. 1, Begon et al., 2005; Clobert, Baguette, et al., 2012; Kendall et al., 1999; Krebs, 2014; Stenseth et al., 2002). The environment may also influence individual state variables (Arrow 2 in Fig. 1), such as sex, body size or body condition which, in turn, also influence demographic rates (Arrow 2-3 sequence in Fig. 1, Charlesworth, 1994; Clobert et al., 2012, 2009; Kooijman, 2010; Le Galliard et al., 2012; Matthysen, 2012; Roff, 2002). Finally, in a feedback loop, changes in demographic rates entail changes in population size and geographic ranges (Arrow 4 in Fig. 1), i.e., changes in the environmental conditions experienced by individuals (e.g., competition, predation, habitat quality, Arrow 5 in Fig. 1).

This feedback loop may be disrupted by global changes, which act at both the environment and individual-state levels. At the environment level, climate change alters temperatures and precipitations (IPCC, 2021). In parallel, a number of human activities, such as harvesting, or their consequences, such as habitat fragmentation, are often selective on sex, physiology, behaviour or body size (Biro and Post, 2008; Cote et al., 2017; Edeline, 2016; Kuparinen and Festa-Bianchet, 2017).

An additional complexity layer comes from state variables being potentially subject not only to plastic changes in response to environmental alterations, but also to evolutionary changes. In particular, emigration from the core habitat, or from the introduction point in the case of biological invasions, selects for increased dispersal propensity through spatial sorting, and is further possibly a competition-avoidance response where poorer competitors disperse from high-density core habitats to low-density marginal habitats (Burton et al., 2010; Chuang and Peterson, 2016; Messager and Olden, 2019; Phillips, 2016; Phillips et al., 2010).

Hence, high dispersers and poor competitors are likely to be overrepresented among immigrants at range edges. Accordingly, results from spread experiments support the prediction that increased dispersal evolves at range edges (Fronhofer and Altermatt, 2015; Ochocki and Miller, 2017; Williams et al., 2016), but evidence for reduced competitive ability is mixed (Duckworth, 2008; Hudina et al., 2015).

Unfortunately, however, the development of process-explicit models of geographic ranges that would integrate these feedbacks is impeded by serious logistical and knowledge limitations. The data needed to estimate demographic rates, as well as their state- and environment-dependency, are hard and costly to acquire in the wild, and are thus typically lacking (Briscoe et al., 2019; Gaston, 2009). Even when present, such data may be poor if detection and/or recapture rates are low, or if population size is small, resulting in inaccurate parameter estimation. A further obstacle to implementing process-explicit models of species ranges is that the eco-evolutionary feedback loops that may operate during range expansion remain a frontier in biology and are poorly understood empirically (Angert et al., 2020; Clobert et al., 2012b; Miller et al., 2020; Travis and Dytham, 2012), which makes it uncertain which mechanisms should be incorporated into models.

In the present study, we propose to circumvent these logistical and knowledge limitations using mark-recapture in outdoor mesocosms of wild-caught, range-core and range-edge animals. Specifically, we quantified the effects of key individual state variables (sex, body size, body condition), climate (temperature), ecology (population density) and geographic range (core vs. edge of an invaded area) on both survival and overland dispersal propensity in red swamp crayfish (*Procambarus clarkii*). This species, well known for dispersing through overland movements (Cruz and Rebelo, 2007; Huner and Barr, 1991; Ramalho and Anastácio, 2015), is considered as one of the most invasive aquatic species worldwide (Oficialdegui et al., 2019; Savini et al., 2010), and poses very serious management problems due to the profound reorganizations they cause in recipient ecosystems (Lodge et al., 2012; Souty-Grosset et al., 2016; Twardochleb et al., 2013). From a taxonomically-wide literature review, we formed detailed predictions for the state, temperature-, density- and range-dependency of *P. clarkii* survival and dispersal (Table 1). Our experimental approach allowed us to fully test all of these predictions.

**Table 1.** Predicted and experimentally-observed in the present study effects of sex, body size, body condition, geographic range (core vs. edge), as well as population density and temperature on both survival and dispersal in the red swamp crayfish (*Procambarus clarkii*) . Detailed statistics for observed effects are provided in Table 2.

| Factor | Relationship to Survival |  |  |  | Relationship to Dispersal |  |  |  |
| --- | --- | --- | --- | --- | --- | --- | --- | --- |
|  | Effect sign |  | Explanation of predicted sign | References | Effect sign |  | Explanation of predicted sign | References |
|  | Predicted | Observed |  |  | Predicted | Observed |  |  |
| Sex male | - | 0 | Males invest less than females in maintenance and future reproduction due to higher hazards | (Kirkwood, 2001; Williams, 1957) | + or - | 0 | Sign depends on relative strengths of opposing selective forces. Male-biased in *Pacifastacus leniusculus* | (Gros et al., 2008; Hudina et al., 2012; Wutz and Geist, 2013) |
| Body size | + | 0 | Larger-bodied individuals dominate competition | (Figler et al. 1999; Edeline and Loeuille 2021 and references therein) | + | + | Sign depends on relative strengths of opposing selective forces. In crayfish, overland and in-stream dispersal is often favoured by a large body size. | (Claussen et al., 2000; Clobert et al., 2009; Kisdi et al., 2012; Moorhouse and MacDonald, 2011; Ramalho and Anastácio, 2015; Thomas et al., 2018; Wutz and Geist, 2013) |
| Body condition | + | - | Higher physical quality increases survival | Multiple evidence, see e.g. Blums et al. (2005) | + or - | + | Sign depends on relative strengths of opposing selective forces. | (Bonte and Peña, 2009; Clobert et al., 2009; Kisdi et al., 2012) |
| High population density | - | - | Increased competition decreases average survival | (Burton et al., 2010; Phillips, 2016; Phillips et al., 2010) see also Appendix 8 for crayfish-specific studies | + | 0 | Competition avoidance favours increased dispersal | (Clobert et al., 2009; Galib et al., 2022) |
| Temperature | - | + | Higher metabolic rate elevates mortality, but sign should depend on temperature relative to thermal optimum | (Angilletta Jr., 2009; Brown et al., 2004; Knies and Kingsolver, 2010; Westhoff and Rosenberger, 2016) | + or - | + | Sign depends on thermal preference | (Angilletta Jr., 2009; Claussen et al., 2000; Le Galliard et al., 2012; Ramalho and Anastácio, 2015) |
| Range edge | - | 0 | A survival-dispersal trade off makes dispersers less able to survive | (Angert et al., 2020; Burton et al., 2010; Messenger and Olden, 2019; Miller et al., 2020; Phillips, 2016; Phillips et al., 2010) | + | 0 | Spatial filtering leads to more dispersive individuals at range edges | (Angert et al., 2020; Burton et al., 2010; Miller et al., 2020; Phillips, 2016; Phillips et al., 2010) |
| High density-by-size interaction | + | + | Benefits from a large body size increase under elevated competition | (Burton et al., 2010; Edeline and Loeuille, 2021; Phillips, 2016; Phillips et al., 2010) | - | 0 | Competition at high density favours dispersal in small, competitively-dominated individuals | (Clobert et al., 2009) |
| Range edge-by-size interaction | - | 0 | Lower competitive ability in Range-edge crayfish make body size less important to survival | (Burton et al., 2010; Edeline and Loeuille, 2021; Messenger and Olden, 2019; Phillips, 2016; Phillips et al., 2010) | - | 0 | Lower competitive ability in Range-edge crayfish makes dispersal less size-dependent | (Clobert et al., 2009; Phillips et al., 2010) |
| Range edge-by-density interaction | - | 0 | Negative effect of high density on survival stronger in less competitive Range-edge crayfish | (Burton et al., 2010; Edeline and Loeuille, 2021; Messenger and Olden, 2019; Phillips, 2016; Phillips et al., 2010) | + | 0 | Lower competitive ability in Range-edge crayfish makes them more sensitive to the positive effect of density | (Clobert et al., 2009; Phillips et al., 2010) |
| Range edge-by-density-by-size interaction | - | 0 | Weaker size-dependency of survival in Range-edge crayfish is more pronounced under high density | (Burton et al., 2010; Edeline and Loeuille, 2021; Phillips, 2016; Phillips et al., 2010) | - | 0 | Weaker size-dependency of dispersal in Range-edge crayfish is more pronounced under high density | (Clobert et al., 2009; Phillips et al., 2010) |

## 2. MATERIALS AND METHODS

### 2.1. Crayfish populations

*P. clarkii* originates from south-central USA (Louisiana and Texas) and was introduced to Europe through both USA-to-Kenya-to-Europe and USA-to-Spain routes (Oficialdegui et al., 2019). The species is highly fecund, tolerant to a large range of environmental conditions (including drought), generally spreads rapidly once introduced, either through secondary human-mediated introductions (Oficialdegui et al., 2020) or through active in-stream and overland dispersal (Cruz and Rebelo, 2007; Tréguier et al., 2018), and often has major impacts on the invaded ecosystems (Souty-Grosset et al., 2016; Twardochleb et al., 2013).

In this study, we used crayfish sampled from natural populations in the Brière area, north-western France, where crayfish belonging to the USA-to-Kenya-to-Europe introduction route were introduced during the early 1980’s (Bélouard et al., 2019; Oficialdegui et al., 2019). Crayfish escaped from a farm and invaded the nearby marsh (90 km^2^), where they now form a large and genetically-homogeneous population (Bélouard et al., 2019). From this range core, dispersers used streams, ditches and, secondarily, overland dispersal to colonize ponds embedded in the surrounding hedgerow landscape, where they founded multiple new populations that may be considered as range-edge populations (Bélouard et al., 2019; Tréguier et al., 2018).

We sampled crayfish from three different range-core locations in the marsh (MB1, MB2 and MB3 as named in Bélouard et al. 2019), hereafter collectively referred to as “Range-core crayfish”, and from three range-edge pond populations genetically differentiated from the marsh source at neutral markers (CB7, I6 and I10 ponds in Bélouard et al. 2019), and hereafter collectively referred to as “Range-edge crayfish”. Further details on the populations used in the experiment are provided in Appendix 1.

In addition to crayfish from the Brière, we sampled crayfish from a third locality, a pond located in Rennes City (about 80 km away from the Brière, Appendix 1), hereafter referred to as an “Outgroup”. These Rennes crayfish have a genetic signature typical of the native range, suggestive of an introduction route that differs from what has been described for the marsh of Brière (Appendix 2). We used these Outgroup crayfish in order to standardize the density treatments in the experiment (see below), and also because we were able to capture Range-edge crayfish only in limited numbers.

### 2.2. Crayfish sampling, marking and maintenance

Crayfish in the Brière were sampled using non-baited traps on May 21^st^, 22^nd^ and 24^th^ 2019. Upon capture, crayfish were determined as male or female from secondary sexual characters (first pleopod pair transformed to gonopods in males), measured for body size (cephalothorax length to the nearest mm, from the tip of the rostrum to the end of the cephalothorax), weighted to the nearest 10^-1^g, and individually marked using 8 mm passive integrated transponders (Biomark HPT8 FDX PIT tags) inserted in the abdominal musculature of the third abdominal segment in the ventral surface. Crayfish were then transported to the PEARL, the INRAE experimental facility for aquatic ecology and ecotoxicology in Rennes (https://www6.rennes.inrae.fr/u3e_eng/), and stocked for 10 days in 8 m^2^ mesocosms under a density of 15 crayfish m^-2^. In stocking mesocosms, crayfish were provided with an excess shelters made from PVC tubes, and were fed with fish food in excess (Le Gouessant, Carpe Extrudée Coul 4). The same procedure was applied to Rennes crayfish, that were PIT-tagged on April 24^th^ and stocked for 40 days.

### 2.3. Experimental mesocosms

Experimental mesocosms were 28, three-m^2^ circular tanks covered with nets to prevent avian predation (Appendix 3A), and equipped with six artificial refuge traps (ART, Green et al. 2018). ARTs consisted in five PVC tubes (250 mm length) glued together and closed by a grid at one end (Appendix 3B), using either 26.5 or 32.5 mm tubes (inner diameter) so as to fit with the needs of both small- and large-bodied crayfish. ARTs sank on the bottom of the mesocosms and provided crayfish with shelters in excess (i.e., one tube provided one shelter, 30 tubes per mesocosm). We initially did not introduce any sediments into the mesocosms, but rapid algal sedimentation provided crayfish with some habitat structure at the bottom of the mesocosms (Appendix 3A).

Each experimental mesocosm was further equipped with an exit trap requiring for trapping that crayfish actively walk out from water, i.e., that crayfish start overland dispersal. Specifically, exit traps were made from a 50L bucket attached to the inner wall of the mesocosm and hanging 30 cm underwater. Trapping required crayfish to climb a ramp made from a plastic net and extending from the bottom of the mescososm up to the rim of the bucket (Appendix 3C). Hence, exit traps captured crayfish that clearly expressed active, out-of-water movements which may be interpreted as the onset of overland dispersal. Crayfish falling into the trap could not escape back to the mesocosm. The bottom of the bucket was pierced with holes to allow water circulation, and PVC tubes provided trapped crayfish with shelters.

Throughout the experiment, crayfish were mildly fed with 418 ± 11 mg (mean ± SD) of fish food per experimental mesocosm per week.

### 2.4. Experimental treatments and recapture

On June 3^rd^ 2019, we started the experiment by transferring crayfish into experimental mesocosms. We introduced randomly-chosen Range-core and Range-edge crayfish at a constant density of 7 crayfish per mesocosm, and varied density using either 2 or 12 Outgroup crayfish per mesocosm, yielding two density treatments (9 or 19 crayfish per mesocosm) and two range treatments (Range-core with Outgroup or Range-edge with Outgroup) in a factorial design (7 replicate mesocosms per treatment, 392 crayfish in total).

Experimental densities (3.0 vs. 6.3 individuals m^-2^) were relatively high compared to naturally observed densities in several crayfish species in invaded areas (including *P. clarkii*), which range from 9.3 10^-3^ to 20.0 individuals m^-2^ (n = 29, mean 2.5, median 1.0, SD 4.7, Bubb et al., 2004; Coignet et al., 2012; Correia & Bandeira, 2004; Galib et al., 2022; Guan, 2000; Lamontagne & Rasmussen, 1993; Pilotto et al., 2008; Wutz & Geist, 2013), and in line with experimentally manipulated densities in enclosures or mesocosms, which range from 0.8 to 14.0 individuals m^-2^ (mean 4.3, median 3.6, SD 3.3, Lodge et al. 1994; Angeler et al. 2003; Rodríguez et al. 2003; Gherardi and Lazzara 2006; Gherardi and Acquistapace 2007; Correia and Anastácio 2008; Matsuzaki et al. 2009; Jackson et al. 2014; Rodríguez-Pérez et al. 2016; Závorka et al. 2020).

At 1-7 days intervals (mean ± SD = 2.4 ± 0.8 days) starting from June 5^th^, both ARTs and exit traps were cleared, recaptured crayfish were recorded for PIT tag number and, so as to proceed rapidly, were returned to their mesocosm without further phenotyping. Two live-recaptured crayfish had lost their tags during the course of the experiment (on 21/06 and 7/08 2019). These were identified based on their sex, size and previous capture histories, and re-tagged. On November 5^th^, after 153 days of experiment, mesocosms were drained and all surviving crayfish were recaptured and recorded for PIT tag number. In total, there were 63 recapture occasions.

All survivors were final-phenotyped following the same protocol as for introductions. Crayfish were anaesthetized and a subsample was dissected for another project, while the remaining ones (n = 231) were frozen at 20°C. Unfortunately, the final-phenotype data was lost due to computer problems, and it was necessary to redo the phenotyping from the remaining, thawed crayfish.

### 2.5. Temperature

Hourly air temperature data for the entire duration of the experiment was obtained from a 5km distant Meteo France weather station (Station météorologique de Rennes-St Jacques, 48°07′N, 1°74′W). Additionally, during 90 % of the duration of the experiment (from June 17^th^ to November 5^th^) we recorded hourly water temperature in six of 28 experimental mesocosms using HOBO Tidbit v2 temperature loggers. A cross-correlation analysis shows that water temperature in the mesocosms was highly correlated with air temperature at the weather station with a 5-hour lag (Appendix 4). For our analyses, we computed mean hourly air temperature during the transition interval between two successive capture occasions.

### 2.6. Multistate capture-recapture model

We estimated the effects of individual- and environment-level factors on probabilities of crayfish survival and overland dispersal using a state-space formulation of a multistate mark-recapture model (Gimenez et al., 2012; Kéry and Schaub, 2012; Lebreton et al., 2009), separating among three states: state A for crayfish present in the mesocosm, state B for crayfish present in the exit trap (i.e., overland dispersal), and Dead. We assumed that crayfish survived in state A between capture occasions *t* and *t*+1, and dispersed to B just prior to *t*+1. Hence, only survival in state A is considered. This assumption was justified by the fact that we did not find any dead crayfish in an exit trap. The state process can be represented by a transition matrix with departure states (time *t*) in rows and arrival states (time *t*+1) in columns:

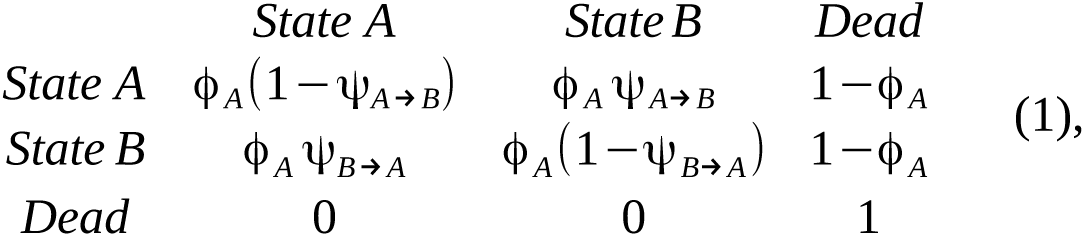

with Φ*_A_* = survival probability, ψ*_A_* _→_ *_B_* = transition probability from state A to state B (i.e., dispersal from mesocosm to exit trap), and ψ*_B_*_→_ *_A_* = transition probability from state B to state A (i.e., no dispersal after having dispersed).

Although all crayfish captured in an exit trap (state B) were in fact effectively returned to their experimental mesocosm (state A), i.e., although the experimental design made state B effectively absent in the rows of the state transition matrix (1), we chose to use this model parametrization so as to separate dispersal performed by previously-dispersing individuals. That is, ψ*_B_*_→_ *_A_* represented probability that crayfish having dispersed to the exit trap did not disperse at next occasion, hereafter referred to as “post-dispersal settlement” probability. Non-symmetric patterns of ψ*_A_* _→_ *_B_* and ψ*_B_*_→_ *_A_* would be suggestive of a cost of dispersal. Note that we also ran a model in which transition probabilities were the same for the “state A” and “state B” lines in the transition matrix (i.e., with effectively no “state B” line) with no change on posterior parameter estimates for ϕ *_A_* and ψ*_A_* _→_ *_B_* .

The observation process is conditional on underlying states, and is described via detection probabilities in each state *p_A_* and *p_B_* . In our experiment, ARTs provided imperfect detection (crayfish present in a mesocosm but not in ARTs when cleared), such that 0< *p_A_* < 1, but crayfish present in exit traps were seen for sure, such that *P_B_* =1 . These observations can be summarized by an observation matrix:

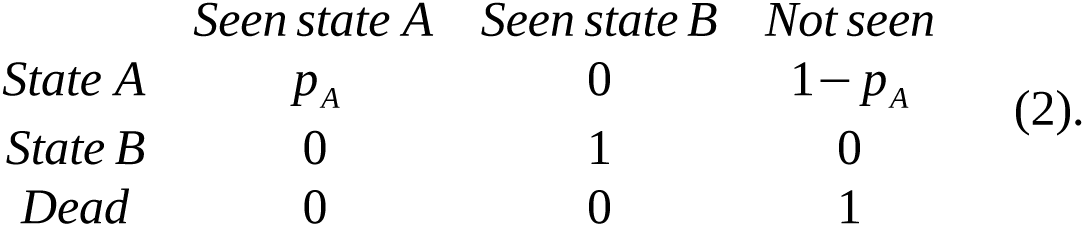

Our model included a “full-detection” formulation with one independent *p_A_* parameter estimated at each occasion *t* for each individual *i*. The model further included three independent Bernoulli GLMMs for the effects of individual- and environment-level factors on probabilities ϕ *_A_*, ψ*_A_* _→_ *_B_*, and ψ*_B_*_→_ *_A_* (generically symbolized by Ω):

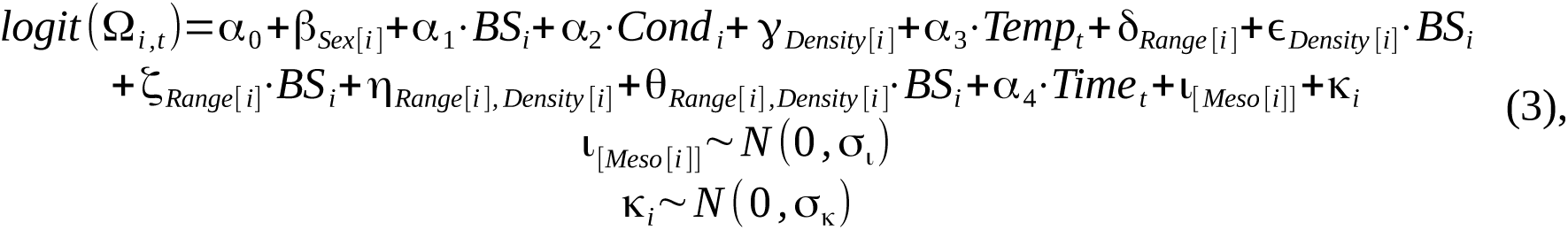

where subscripts *i* and *t* stand for crayfish individuals and capture occasions, respectively, α_0_ is model intercept (see below), *Sex* is a two-level factor (Male or Female), *Range* has three levels (Range-core, Range-edge, Outgroup), *Density* has two levels (9 or 19 crayfish per mesocosm), *Temp* is the mean of hourly air temperature during the transition interval between two capture occasions, *Time* is duration of the transition interval in days, and *Meso* is mesocosm identity (numbered from 1 to 28).

In Eq. 3, *BS* is initial body size at introduction (cephalothorax length, mm), and *Cond* is initital individual body condition measured as the residual from the ln(Mass_i_) / ln(*BS*_i_) linear regression. Our model thus assumes that initial body size and condition differences among individuals were retained during the whole course of the experiment. Accordingly, the subset of surviving crayfish that was phenotyped after thawing at the end of the experiment (see above) showed that initial body sizes and masses were very good predictors of final lengths and masses, i.e., validated our assumption. Specifically, the length regression had a R^2^ > 0.99 and slope = 1.07 indicating 7 % average growth in length in survivors, and the mass regression had R^2^ > 0.94 and slope = 1.17 indicating 17 % average growth in mass in survivors. The lower R^2^ for the mass regression might be partly explained by the freezing-thawing sequence, that may generate random water losses.

The ι and κ random effects in Eq. 3 accounted for mesocosm and individual (overdispersion) effects. Note that ϕ *_A_* was not overdispersed and, hence, its GLMM did not include a κ parameter. Eq. 3 was fitted using an “effect” parametrization as is the default in R, such that the intercept α_0_ corresponds to mean response in Range-core females at low density. To evaluate the main effects of covariates, we also fitted a version of Eq. 3 with no interaction term.

To test for nonlinearities in the effects of temperature on crayfish survival and dispersal, we also fitted a version of the model in which Eq. 3 included a *Density*-by-*Temp*, a *BS*-by-*Temp* and a *Density*-by-*BS*-by-*Temp* interactions. None of these interactions were statistically significant, and we thus discarded this version of the model from our subsequent analyses.

We fitted the model using Markov chain Monte Carlo (MCMC) in JAGS (Plummer, 2003) through the jagsUI package (Kellner, 2019) for R 4.2.1 (R Core Team, 2022). All numeric covariates were centred to zero mean and 0.5 standard deviation as recommended by Gelman (2008) for logistic regressions. We used vague priors on all parameter distributions. Specifically, we chose Cauchy robust priors with scale 2.5 for regression parameters (Gelman et al., 2008), uniform distributions on the [0,1] interval for probabilities, and uniform distributions on the [0,5] interval for variance parameters. Additionally, to test posterior sensitivity to the priors, we also ran the model using as priors uniform distributions on the [-10,10] interval for regression parameters in Eq. 3.

We ran three parallel MCMC chains for 10000 iterations (burnin of 2000 iterations) thinned at a 4 iteration interval. We assessed parameter convergence using the Gelman-Rubin statistic which compares the within- to the between-variability of chains started at different and dispersed initial values (Gelman and Rubin, 1992). We tested for the significance of effects in (Eq. 3) through computing MCMC p-values as twice the proportion of the posterior which sign was opposite to the sign of the posterior mode. We further evaluated the identifiability of model parameters by computing the overlap between prior and posterior distributions using the BEST package in R (Kruschke and Meredith, 2021). A high prior-posterior overlap (PPO) indicates that information from the data is weak, such that the posterior parameter distribution is strongly dependent on the prior distribution. A PPO greater than 35 % is considered as indicative of a weak parameter identifiability (Gimenez et al., 2009).

## 3. RESULTS

### 3.1. Phenotypes

Crayfish had a balanced sex ratio (51 % females, see Appendix 1 for population-specific numbers). Cephalothorax length ranged from 24.9 to 66.2 mm (mean ± SD = 44.7 ± 6.8 mm), and was similar among Range-core (42.7 ± 7.1 mm) and Range-edge crayfish (42.2 ± 6.4 mm), but was significantly larger in Outgroup crayfish (46.9 ± 6.2 mm, Appendix 1; estimate = 4.22, ddl = 392, res. ddl = 389, t-value = 5.01, p-value < 0.001). Outgroup crayfish also had a significantly higher body condition than Brière crayfish (estimate = 0.10, ddl = 392, res. ddl = 389, t-value = 7.31, p-value < 0.001).

### 3.2. Temperature

Mean air temperature during a recapture interval ranged from 9.3 to 26.9°C (Fig. 2). Maximum air temperature was 39.4°C. There was a high positive correlation between mean and maximum temperature (cor = 0.93) and between mean and variance temperature (cor = 0.68, data not shown).

**Fig. 2.**
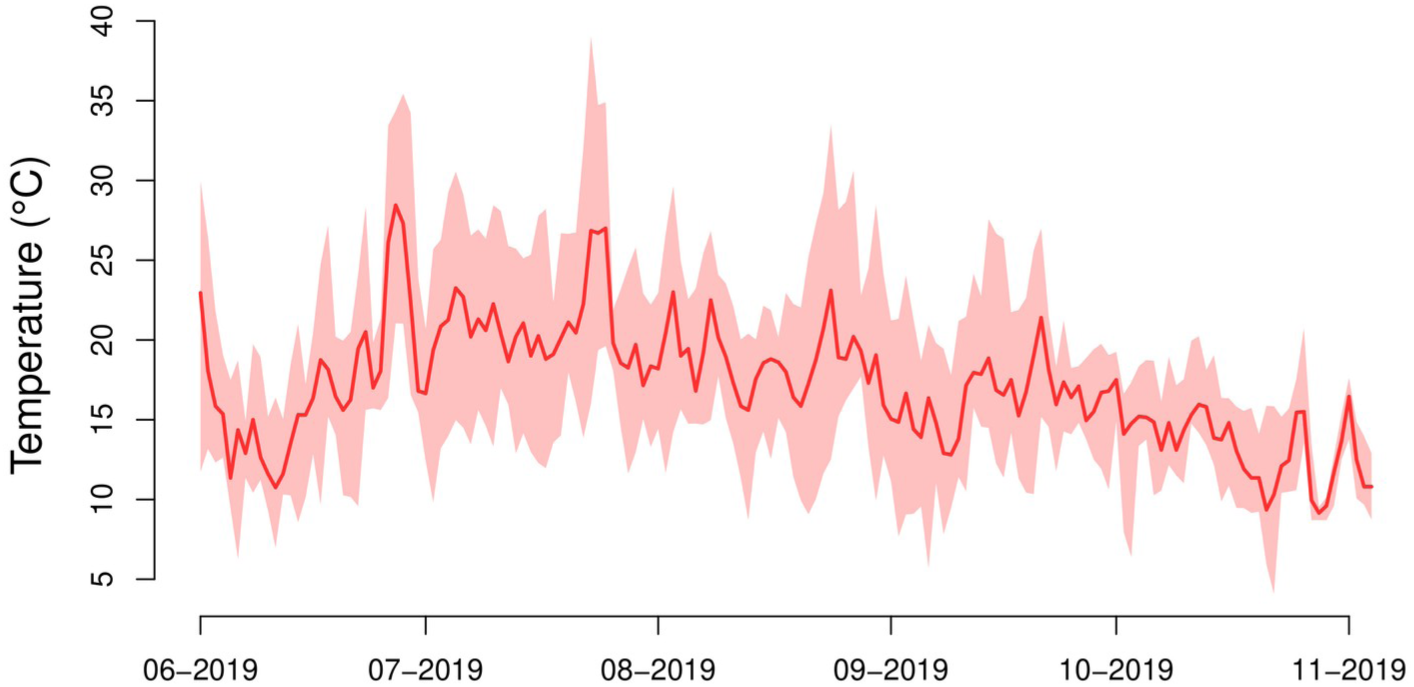
Daily air temperature during the experiment. Solid line: median daily temperature. Ribbon: 95 % observed interval.

### 3.3. Survival

Posteriors for regression parameters in Eq. 3 were insensitive to the choice of Cauchy or uniform priors (compare Table 2 with Appendix 5). Prior-posterior overlaps (PPOs) for survival parameters were below 35 %, except for η[Range edge, HD] (PPO = 36 %), indicating no major identifiability problem (Appendix 6A). In particular, none of the posteriors associated with significant effects were weakly identifiable.

**Table 2.**
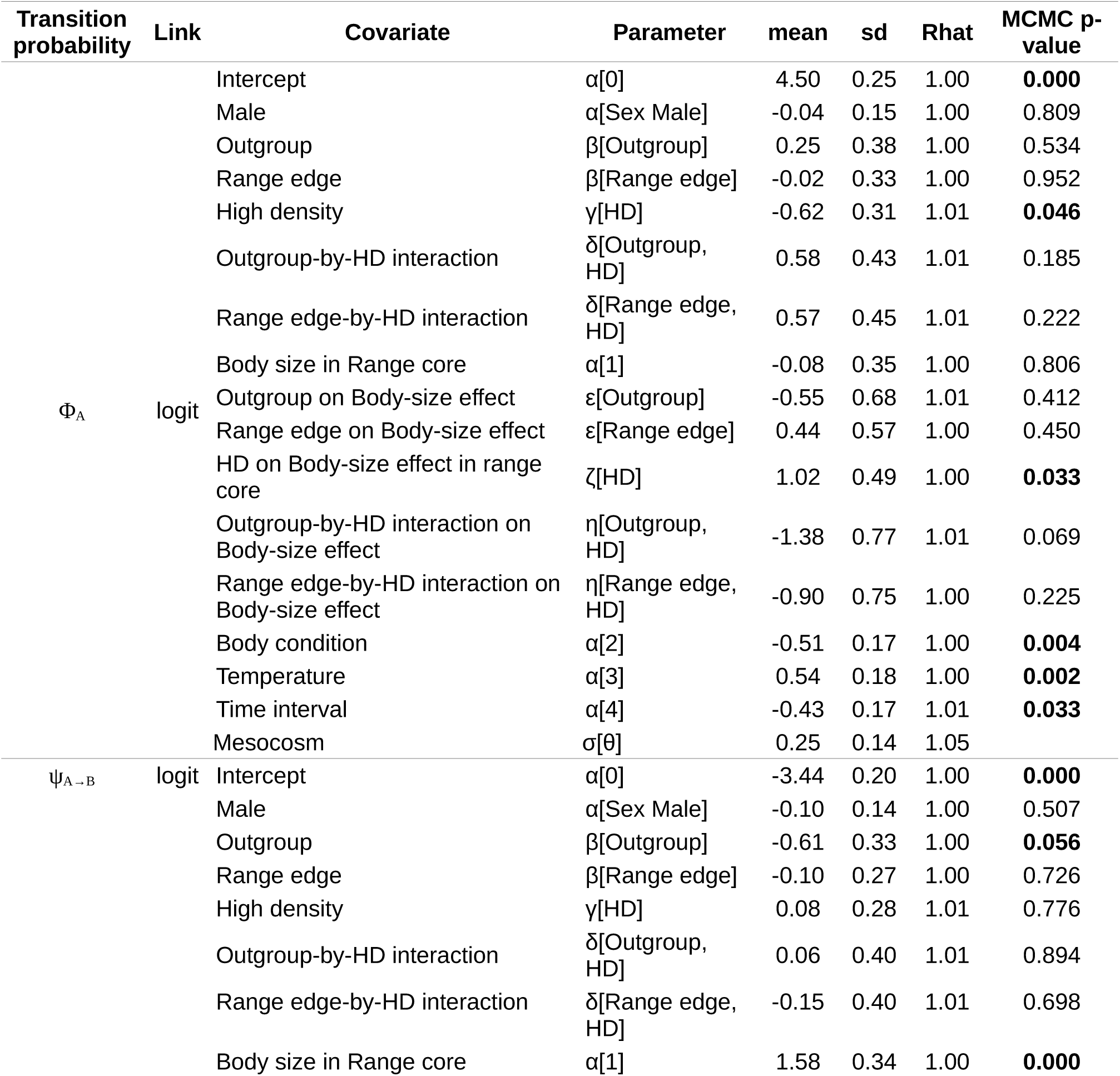
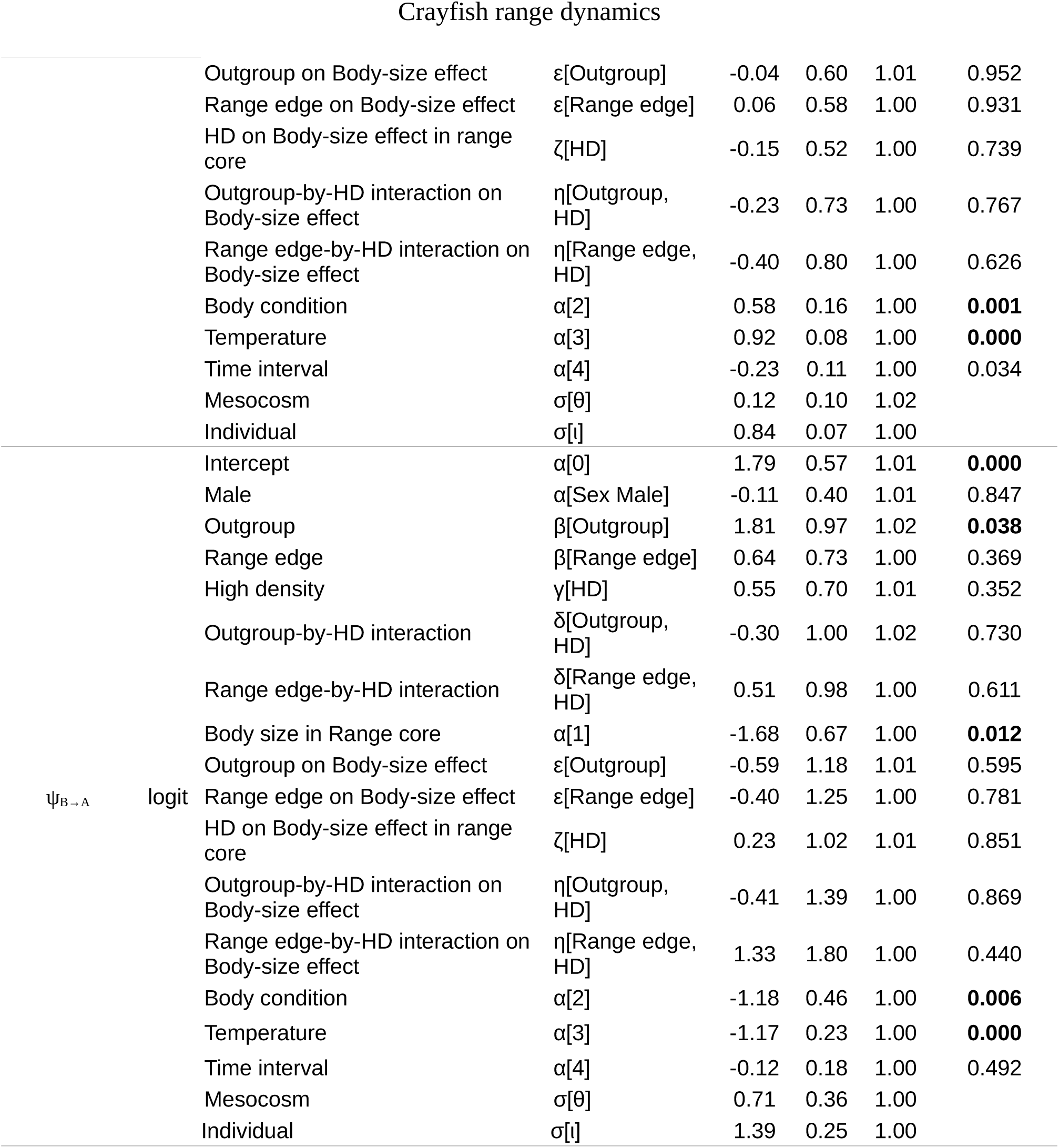
Parameter estimates from the multistate mark-recapture model. Φ_A_: survival probability, ψ_A→B_: dispersal probability, ψ_B→A_: post-dispersal settlement probability. The model is described in Eq. 3 and was fitted using an “effect” parametrization as the default in R. MCMC p-values were computed as twice the proportion of the posterior which sign was opposite to the sign of the posterior mode. P-values significant at the 5 % risk are bold faced. MCMC p-values are not relevant for variance parameters (random effects) that are constrained to be non-zero. HD: High density, LD: Low density. Parameter estimates for a model including only main effects in Eq. 3 (i.e., no interaction) are provided in Appendix 7. NB: The survival model for Φ_A_ was not overdispersed and did not include an “individual” random effect. Quantitative covariates were standardized to zero mean and 0.5 standard deviation.

Crayfish survival probability during a time interval ranged from 0.9 to 1 (Figs. 3a-d). These high probabilities were due to the short duration of a time interval (2.4 days on average). Over the entire duration of experiment (153 days), 51.5 % of crayfish survived.

**Fig. 3.**
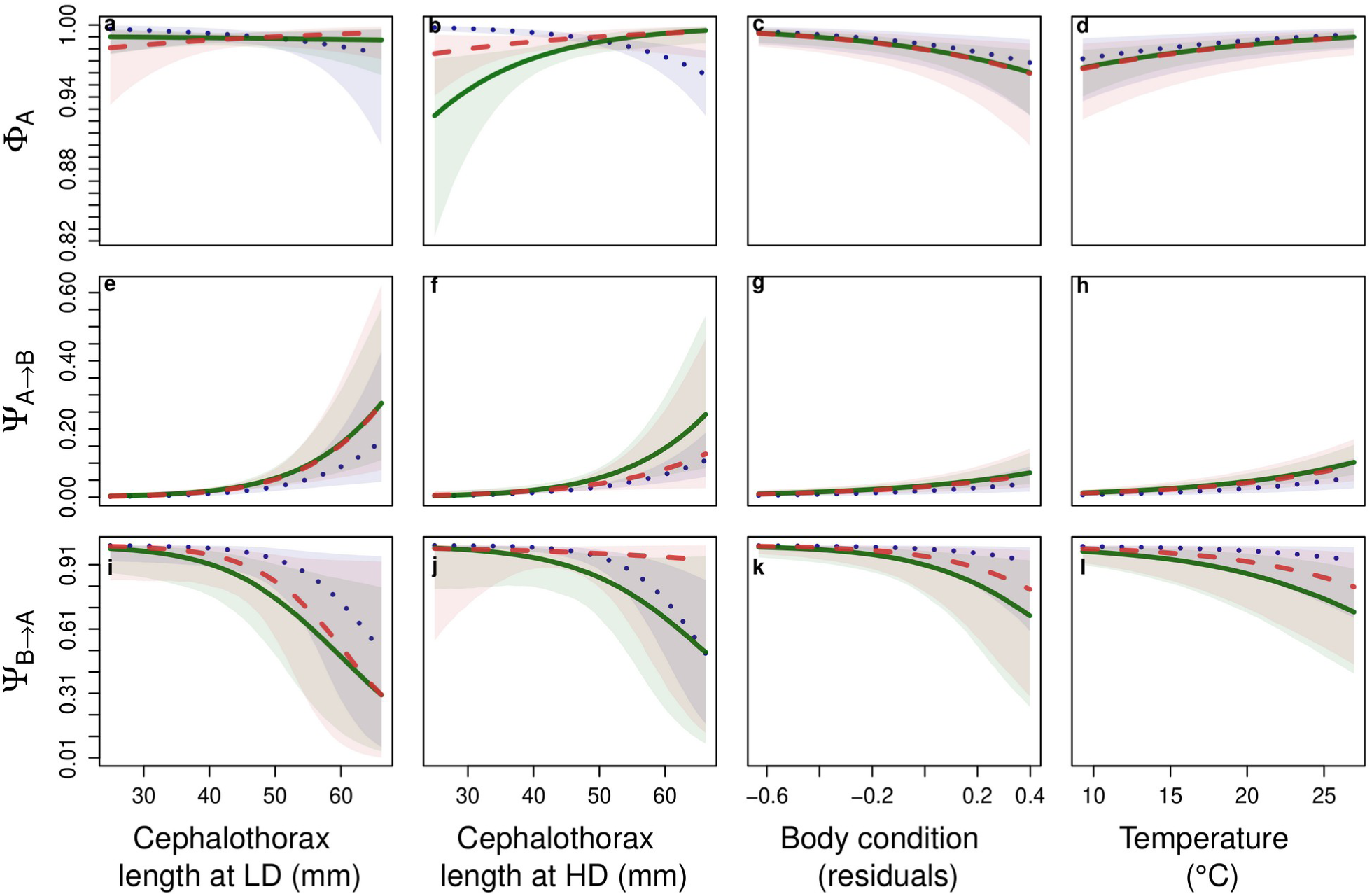
Predicted effects of body size-by-density interaction, body condition and temperature on crayfish survival probability Φ_A_, dispersal probability Ψ *_A →B_* and post-dispersal settlement probability Ψ*_B → A_*. Lines represent median prediction, ribbons indicate 95 % credibility intervals. Green, solid lines: Range-core crayfish. Red, dashed lines: Range-edge crayfish. Blue, dotted lines: Outgroup crayfish. LD and HD stand for low and high crayfish density, respectively. All probabilities are for a transition interval of 2.4 ± 0.8 days (mean ± SD). Predictions were made from parameters estimates of Eq. 3 by setting the Sex factor to female crayfish, and predictions for the effects of body condition and temperature were further made by setting the Density factor to HD. Credible intervals are not displayed for clarity reasons.

Larger-bodied crayfish had increased survival probability under high (but not low) density conditions, a trend mostly present in Range-core crayfish (Figs. 3b *vs*. 3a, α[1] *vs*. ε[HD] parameters in Table 2). Specifically, average survival probability during a 2.4-day time interval was 9.57 10^-1^ in the smallest crayfish (24.9 mm cephalothorax length) and 9.98 10^-1^ in the largest crayfish (66.2 mm, probabilities predicted for Range-core female crayfish under high density, all other quantitative covariates fixed to their average value).

Higher crayfish density (Figs. 3b vs. 3a, γ[HD] parameter in Table 2) and higher individual body condition (Fig. 3c, α[2] parameter in Table 2) both decreased crayfish survival probability. Note that the negative effect of density on crayfish survival was weaker and not significant when evaluated as a main effect in a model that did not include any interaction (Appendix 7).

Warmer temperatures had a positive effect on crayfish survival probability (Fig. 3d, α[3] parameter in Table 2). When all other quantitative covariates were fixed to their average value, raising mean temperature from its minimum (9.3°C) to its maximum (26.9°C) increased mean survival probability during a 2.4-day time interval from 9.81 10^-1^ to 9.95 10^-1^ (predictions computed for an average-sized, Range-core female at low density).

There was no significant effect of the geographic range on crayfish survival probability (δ[Range edge] parameter in Table 2), nor there was any significant range-by-body size (ζ[Range edge] parameter in Table 2), range-by-density (η[Range edge, HD] parameter in Table 2, a weakly identifiable parameter) or range-by-body size-by-density interaction (θ[Range edge, HD] parameter in Table 2). Note that we also tested for range-by-temperature and range-by-temperature-by-density interactions and did not find any significant interaction (results not shown).

### 3.4. Dispersal

PPOs were below 35 % for all dispersal parameters except for η[Outgroup, HD] (PPO = 37 %) and for η[Range edge, HD] (PPO = 40 %), indicating that the identifiability of our model parameters was generally satisfying (Appendix 6B). The probability for an individual crayfish to disperse to an exit trap during a time interval ranged from 0 to 0.3 (Figs. 3 e-h), while post-dispersal settlement probability ranged from 0.3 to 1 (Figs. 3 i-l). Hence, crayfish were mainly sedentary during our experiment.

Dispersal probability was almost 0 in crayfish smaller than 45 mm in cephalothorax length, but increased up to 0.1-0.3 in the largest-bodied individuals, a significant body-size effect (Figs. 3e and 3f, α[1] parameter in Table 2). Specifically, mean dispersal probability increased from 3.07 10^-3^ in a small-bodied crayfish (25 mm cephalothorax length) to 2.86 10^-1^ in a large-bodied crayfish (66 mm, predictions computed for a Range-core female at low density, all other covariates were fixed to their average value).

Increasing body condition, as well as warmer temperatures, also had significantly positive effects on crayfish dispersal probability (Figs. 3g and 3h, respectively, α[2] and α[3] parameters in Table 2). When all other quantitative covariates were fixed to their average value, raising mean temperature from its minimum (9.3°C) to its maximum (26.9°C) increased mean dispersal probability during a 2.4-day time interval from 1.17 10^-2^ to 9.46 10^-2^ (predictions computed for an average-sized, Range-core female at low density)

Neither crayfish density nor geographic range did influence crayfish dispersal probability, either directly or in interaction with body size (Figs 3e and 3f, Table 2).

The identifiability of model parameters for post-dispersal settlement probability was low, since 9 over 16 parameters had prior-posterior overlaps exceeding 35 % (Appendix 6C). However, the specific response of post-dispersal settlement to major covariates (body size, body condition and temperature, as displayed in Fig. 3) was identifiable (α[1], α[2] and α[3] parameters, PPOs ranging from 13 to 27 %, Appendix 6C), and seemed generally symmetric to the response of dispersal probability (Figs. 3i-l vs 3e-h, Table 2), suggesting that dispersing at time *t* did not alter dispersal probability at time *t*+1 with respect to covariates.

## 4. DISCUSSION

Our study shows in the red swamp crayfish that survival and dispersal are under the control of both individual- and environment-level factors (Fig. 1). Below, we first discuss these results separately for each set of factors, and then we discuss the benefits of our findings for range-dynamics modelling and applied issues with invasive *P. clarkii*.

### 4.1. Individual state variables

As predicted (Table 1), a larger body size increased crayfish survival probability, but this effect was significant only under a high density and in Range-core crayfish. This result highlights density dependent selection for a larger body size in *P. clarkii*, as previously reported in other taxa (collembolans, fish and lizards), and presumably resulting from a dominance of large-bodied individuals in interference competition and cannibalism (see Edeline and Loeuille 2021 and references therein). Accordingly, larger-bodied crayfish dominate in competition for shelters (Figler et al., 1999; Rabeni, 1985; Ranta and Lindström, 1993), and crayfish with larger chela have a higher probability of winning agonistic encounters (Graham and Angilletta, 2020).

A larger body size further had a strong positive effect on crayfish dispersal probability in our experiment, a finding that is in line with a previous report of overland movements being predominantly performed by sexually mature (i.e., adult) individuals in *P. clarkii* (Ramalho and Anastácio, 2015). In *Pacifastacus leniusculus,* both overland and in-stream dispersal also increase with increasing body size (Bubb et al., 2004; Moorhouse and MacDonald, 2011; Thomas et al., 2018; Wutz and Geist, 2013), suggesting that a larger body size elevates a general propensity for movements, and might represent a major component of dispersal syndromes in crayfish.

This strong and taxonomically-conserved positive effect of body size on crayfish movements is probably a response to multiple convergent selective pressures, in link with physiological and ecological constraints. A larger body size decreases the mass-specific energetic cost of transport of both walking or running animals (Peters, 1983), and further confers a higher body volume-to-surface ratio which increases resistance to dessication during overland movements (Antoł et al., 2021; Claussen et al., 2000). Additionally, walking speed also increases with body size (Claussen et al., 2000), and larger-bodied dispersers thus have increased probability to find a water body before dessication. Finally, larger-bodied individuals dominate in interference competition for food and contests for shelters (see above), and thus probably also enjoy reduced costs of settlement during the colonization of novel habitats. Hence, larger-bodied crayfish thus probably enjoy a strongly decreased cost-to-benefit ratio of dispersal.

In accordance with the general expectation that increased energy stores decrease the survival cost of long-distance movements (Table 1), we found that body condition was positively linked to dispersal probability in crayfish. However, in contrast with our predictions we found that a higher body condition was also associated with a lower survival probability. In shrimps, a higher nutritional status is associated with increased moulting probability (Lemos and Weissman, 2021; Sharawy et al., 2019). Maybe, in crayfish also a higher body condition was possibly indicative of a higher energetic status and of a higher probability of moulting, which entails increased mortality in crayfish (Taugbøl and Skurdal, 1992). In particular, newly-moulted crayfish have a soft shell and are strongly exposed to cannibalism.

Also contrary to our expectations (Table 1), survival and dispersal were both sex-independent. This result suggests that, in *P. clarkii*, hazards and costs of dispersal are similar among males and females. In *P*. *leniusculus*, dispersal was reported to be sometimes sex-independent (Bubb et al., 2004; Galib et al., 2022), and sometimes male-biased (Wutz and Geist, 2013), which might suggest that, in crayfish in general, the sex-dependency of dispersal is context-dependent and versatile.

Our predictions were also refuted regarding the effects of spatial sorting on competitive ability and dispersal propensity. Range-core and Range-edge crayfish did not differ in either survival or dispersal probabilities, nor they did in their response to crayfish density variation. Such an absence of any effect of the geographic range contrasts with theoretical expectations (Table 1), and is maybe surprising given the large genetic divergence of Range-edge crayfish at neutral markers (Bélouard et al., 2019), but agrees with previous results showing no divergence among Range-core and Range-edge crayfish populations in the Brière for the fluctuating asymmetry of morphological traits (Bélouard et al., 2019). This apparent paradox may result from both genetic and selective processes.

Without selection, divergence in quantitative traits should equal the divergence at neutral marker loci (Leinonen et al., 2013, 2008). However, this equality does not hold for complex traits such as dispersal syndromes (Mackay et al., 2009; Saastamoinen et al., 2018), because allelic interactions within (dominance) or between (epistasis) loci reduce the divergence at quantitative traits compared to the neutral expectation (Goudet and Martin, 2007; Whitlock, 2008). Additionally, density dependent selection in edge habitats may have rapidly erased the effects of spatial sorting to make Range-edge crayfish similarly dispersive and competitive as their Range-core counterparts. Strong density-dependent selection is particularly likely to occur in the “boom” phase of large demographic increase that sometimes immediately follows a colonization event, due to colonists arriving in an empty ecological niche with lots of resources available (Strayer et al., 2017). Which of these different mechanisms best explains phenotypic similarity among Range-core and Range-edge crayfish in the Brière requires further investigation.

### 4.2. Environmental variables

As expected (Table 1), crayfish density in mesocosms had a marginally-significant negative effect on the survival of Range-core crayfish. However, this effect was weaker than expected since it was not significant when evaluated simultaneously across all crayfish origins or sex (main-effects model, γ[High density] parameter, Appendix 7). This weakness probably reflects limited density variation (from 3.0 to 6.3 individuals m^-2^) in presence of an excess shelters (30 shelters for a maximum of 19 crayfish introduced per mesocosm). Accordingly, an average relationship between survival probability and stocking density for different experiment durations, as reconstructed from literature data (Appendix 8), predicts largely overlapping survival probabilities at these two densities and for an average time interval of 153 days corresponding to the duration of our experiment (0.64-0.85 at 3.0 individuals m^-2^ and 0.58-0.80 at 6.3 individuals m^-2^). Hence, we conclude that the density variation we applied in our experiment, although located in the upper range of naturally-observed densities, entailed relatively mild changes in competition and, hence, resulted in weak effects on survival and no effect on dispersal.

Finally, temperature was the only environmental factor that had consistent effects on crayfish demographic rates during our experiment: both survival and dispersal probabilities increased with increasing temperature. All biological rates are expectedly maximized at intermediate temperatures (Table 1), which correspond to a specific thermal optimum that may vary among species (Angilletta Jr., 2009; Knies and Kingsolver, 2010). Crayfish are no exception and reach maximal rates of somatic growth (Westhoff and Rosenberger, 2016) and dispersal (Claussen et al., 2000; Ramalho and Anastácio, 2015) at intermediate temperatures. However, when biological rates are measured over a narrow temperature range, their hump-shaped relationship to temperature may not be observed, and authors report either a positive or negative relationship. Accordingly, warmer temperatures are reported to either increase or decrease rates of crayfish survival rate (Harlıoğlu, 2009; Verhoef and Austin, 1999; Webster et al., 2004). In *P. clarkii*, somatic growth rate, which is a putative surrogate for survival rate, is maximal in the 22-30°C temperature range (Westhoff and Rosenberger, 2016), while overland dispersal seems maximal at temperatures ranging from 16 to 24°C (Ramalho and Anastácio, 2015). These temperatures will more often be met in the future, as hot days will get hotter and more frequent (IPCC, 2021).

In Europe for instance, climate projections suggest that, by 2081-2100 and compared to the 1995-2014 period, temperatures will increase by 1.25 to 2.65°C under the SSP1-2.6 scenario (low GHG emissions), and by 2.45 to 4.85°C under the SSP5-8.5 scenario (high GHG emissions, IPCC, 2021, page 14). Assuming an annual average reference temperature equal to 14.3°C, the mean summer (May-September) temperature in Rennes City^1^, these two warming scenarios are predicted by our model to result in a 1.5 to 5.2 % increase in *P. clarkii* monthly survival probability, and in a 14 to 44 % decrease in the number of days needed for *P. clarkii* to reach an overland dispersal probability equal to 1 (Appendix 9). Hence, although care should be taken in extrapolating experimental results to the real world, our results suggest that climate warming will potentially favour a significant expansion of the crayfish geographic range.

### 4.3. Implications for the projection and management of geographic ranges

Correlative species distribution models based on climate envelopes also predict a geographic range expansion of *P. clarkii* under projected climate change scenarios (Capinha et al., 2013; Liu et al., 2011; Zhang et al., 2020). Our results are consistent with this prediction and, in addition, provide a mechanistic, demography-based understanding of the underlying processes, that may serve as a starting point to building process-explicit models of range dynamics for *P. clarkii* (Fig. 4).

**Fig. 4.**
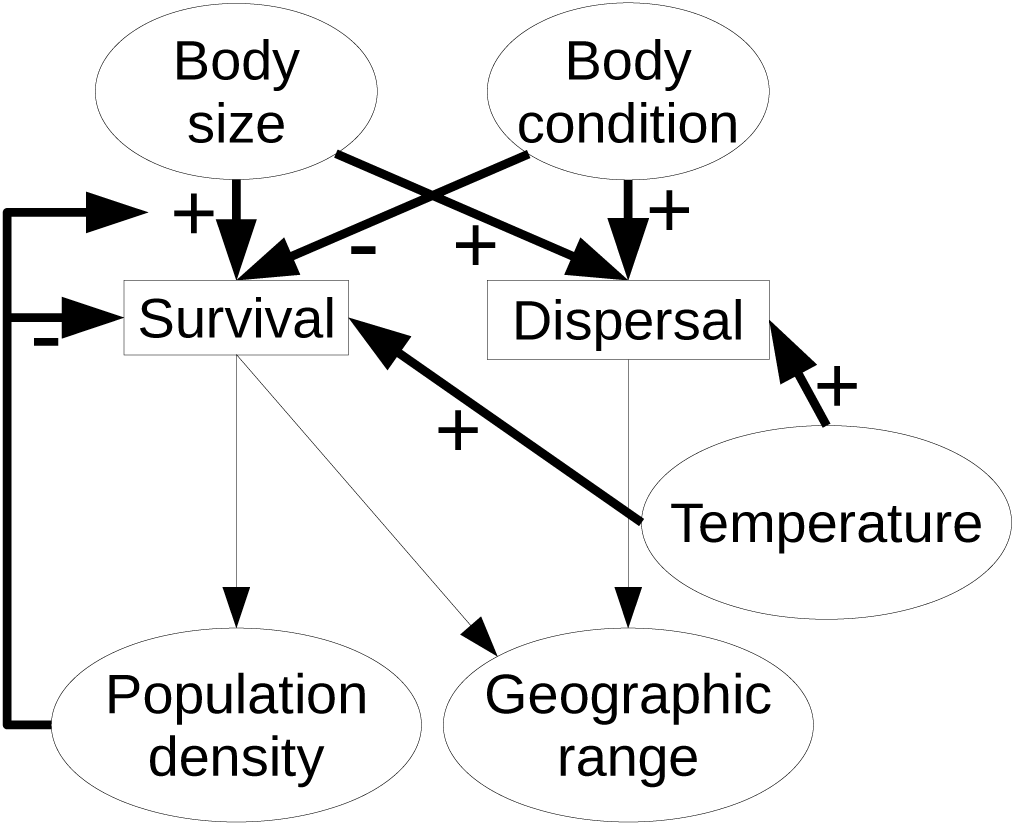
Adaptation of the conceptual framework in Fig. 1 to *P. clarkii* range dynamics. Increased survival and dispersal are assumed to both increase the geographic range. Bold arrows indicate causality links that were investigated in the present study. The arrow pointing from population density to the positive effect of body size on survival indicates that this positive effect was present only under high crayfish density.

In the context of a spreading biological invasion, improvement of containment strategies appears as a crucial management target. Our results suggest that crayfish containment could be based on manipulating crayfish body size. Specifically, mechanically culling large-bodied crayfish through, e.g., selective trapping, might promote an immediate crayfish containment through reducing both average survival and dispersal probabilities in the population (Fig. 4). Culling-based containment might become even more efficient over the long term because, after several generations, selective removal of large-bodied individuals induces an evolutionary change towards smaller body sizes and earlier maturation, which would not only cause crayfish to disperse less, but are also traits generally associated with lower population growth rates (Evangelista et al., 2021, 2020; Heino et al., 2015; Hutchings, 2005). Such harvest-induced evolutionary changes are often considered as slowly reversible (Law, 2000), which is desirable from a long-term management perspective.

However, whether or not long-term culling of large-bodied individuals represents an efficient containment strategy remains to be effectively tested in the field. On the short term, removal of large-bodied individuals relaxes competition and favours a plastic increase in rates of somatic growth and reproduction of survivors, which led some authors to conclude that culling is inefficient at controlling crayfish populations (Gherardi et al., 2011). Additionally, locally-lower population densities may possibly trigger increased immigration from surrounding, non-culled areas where densities are higher (Moorhouse and MacDonald, 2011). Hence, selective culling of large-bodied crayfish might be better envisioned as a large-scale management strategy which, after an initially strong culling effort, may provide good results if milder removal may be sustained over temporal and spatial scales encompassing multiple crayfish generations across the whole (meta)population range.

### 4.4. Conclusions

Our study demonstrates the power of individual mark-recapture in mesocosms to disentangle climate, ecological and evolutionary drivers of demographic rates. Our results show that survival and overland dispersal in *P. clarkii* are under the control of both climatic and ecological factors that act at both the individual and population levels, but suggest no rapid trait evolution at range edges. Future studies should extend the approach to other demographic rates, i.e., somatic-growth and maturation rates, as well as to measuring their dependency to a gradient of environmental conditions. Complementary research is also needed to include other means of dispersal, because *P. clarkii* actively disperses not only overland or through waterways (Cruz and Rebelo, 2007; Tréguier et al., 2018), but also passively, as evidenced by the role played by humans or animals in their current geographic distribution (Acevedo-Limón et al., 2020; Anastácio et al., 2014; Capinha et al., 2013). Finally, future comprehensive mark-recapture experimental data could be coupled to more readily-available count data from natural populations in integrated population models (Briscoe et al., 2019; Pagel and Schurr, 2012; Schaub and Kéry, 2021). This new class of models will provide more robust estimates of demographic rates under a wider range of ecological and climatic scenarios and, we hope, will represent high-class tools for the management of established invasive crayfish populations under global change.

## Acknowledgements

We are grateful to Flavie Perrier and Owen Nino for help with collecting crayfish in the marsh of Brière, and to Mattéo Le Goues for help with the genetic analysis of crayfish from Rennes. Permission for crayfish capture and transport were provided by the Préfecture de la Loire-Atlantique (license number 2019/SEE-53), and authorization for crayfish retention was granted to U3E by the Direction Départementale de la Cohésion Sociale et de la Protection des Populations d’Ille et Vilaine.

## Author contribution

E.E., J-M.P. and E.P. designed the experiment and captured and marked the Brière crayfish. Y.B. and A.S. collected the recapture and final phenotype data, E.E. analysed the data and led manuscript writing. All authors participated in manuscript editing and revisions.

## APPENDICES FOR

## APPENDIX 1. Details of the crayfish populations used in the experiment

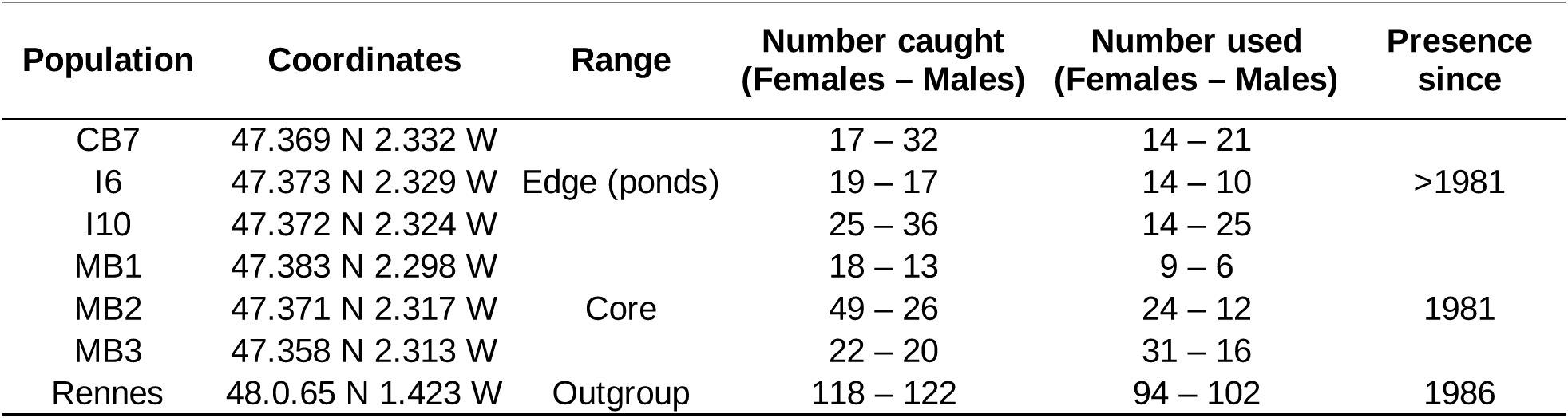

APPENDIX 1. Details of the crayfish populations used in the experiment.
| Population | Coordinates | Range | Number caught<br>(Females – Males) | Number used<br>(Females – Males) | Presence<br>since |
| --- | --- | --- | --- | --- | --- |
| CB7 | 47.369 N 2.332 W |  | 17 – 32 | 14 – 21 |  |
| I6 | 47.373 N 2.329 W | Edge (ponds) | 19 – 17 | 14 – 10 | >1981 |
| I10 | 47.372 N 2.324 W |  | 25 – 36 | 14 – 25 |  |
| MB1 | 47.383 N 2.298 W |  | 18 – 13 | 9 – 6 |  |
| MB2 | 47.371 N 2.317 W | Core | 49 – 26 | 24 – 12 | 1981 |
| MB3 | 47.358 N 2.313 W |  | 22 – 20 | 31 – 16 |  |
| Rennes | 48.065 N 1.423 W | Outgroup | 118 – 122 | 94 – 102 | 1986 |

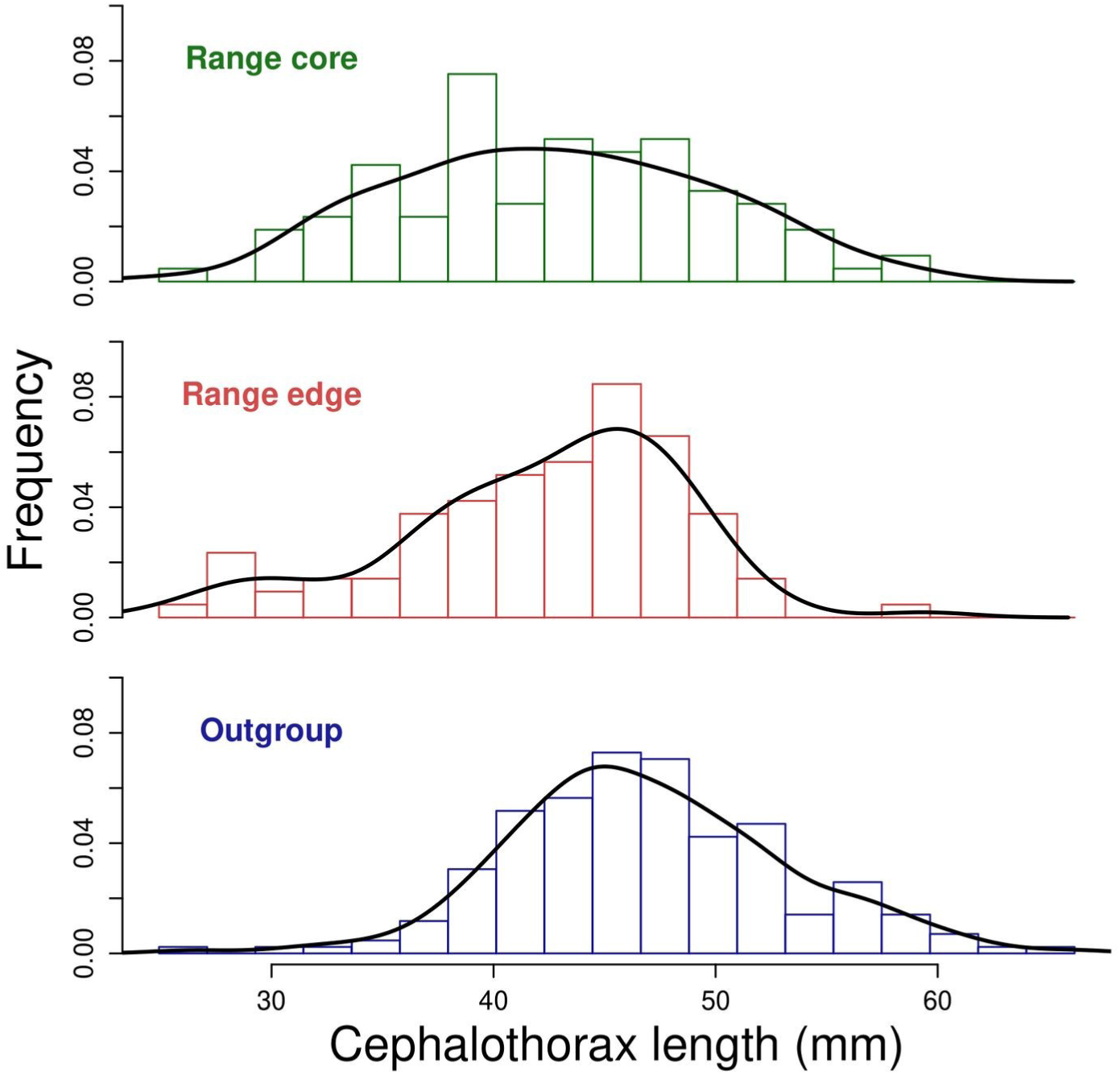

## APPENDIX 2. Genetics of the Rennes (“Outgroup”) crayfish population.

### Aim

—The invasion history of the pond from which our Outgroup individuals originate was unknown. We tested the hypothesis that this invasion followed the same introduction route as red swamp crayfish (*Procambarus clarkii*) from Brière, namely, the USA-Kenya introduction route (Oficialdegui et al., 2019). Besides looking for the origin of this specific pond, we also tested new ways for obtaining DNA material from crayfish. In that aim, we collected three kinds of samples on all individuals: we sampled with sterile cotton swabs (i) the buccal region and (ii) haemolymph leaking out of the abdomen while PIT-tagging. We also collected (iii) muscle from the abdomen when individuals were sacrificed as a classical way to obtain genomic DNA.

### Methods

—30 individuals were collected and stocked in an outdoor mesocosm at the PEARL (https://www6.rennes.inrae.fr/u3e_eng/). Swabs were collected on all individuals at two capture occasions to check whether results were reproducible. At the first capture event, individuals were marked with Biomark HTP8 PIT-tags and leaking haemolymph was collected with a cotton swab. At the second capture event three days later, crayfish were jabbed again with a PIT-tag syringe and a second haemolymph swab was collected, then individuals were sacrificed and their muscle was sampled.

DNA was extracted from all (n=150) samples with the NucleoSpin Tissue kit (Macherey-Nagel), following manufacturer’s instructions. This protocol was slightly modified for swabs, for which we doubled buffer volumes: we added 360µl of the initial (T1) lysis buffer and 25µl of proteinase K to the swab, and then used 400µl of buffer B3 and 420µl of ethanol in the following steps. All extracts were quantified using a DS-11 spectrophotometer (DeNovix).

Because at least one of our sampling protocol targeted extra-corporal DNA (buccal swabs), which is potentially degraded, we chose to amplify and sequence a short mitochondrial DNA fragment (Broquet et al., 2007). Primers were designed from the alignment of published CO1 sequences: *P. clarkii* haplotypes from Oficialdegui et al. (2019) (NCBI accession numbers MK026671 to MK026718) and six CO1 sequences taken from other crayfish species (species names and NCBI accession numbers: *Procambarus virginalis*, LR884234.1; *Procambarus fallax*, LC228303.1; *Procambarus riojae*, KX238226.1; *Procambarus hoffmanni*, KX238193.1; *Procambarus geminus*, JX514457.1; *Pacifastacus leniusculus*, MK439898.1). The resulting primers, PC-FCO1202 (5’-GGGGTATAGTTGAGAGAGGAGT -3’) and PC-RCO1202 (5’-GGTATTCGATCCATGGTTATCCC -3’), amplify a 202bp CO1 fragment in *P. clarkii*. PCR amplifications were carried out in a total volume of 50 µl containing 1µM of each primer, 1mM MgCL2, 1X GoTaq G2 (Promega) PCR Mix, 1.25 units of GoTaq G2 (Promega) Taq Polymerase, 2µl DNA, the whole being completed with sterilized water. DNA samples obtained from muscles were diluted 10 times to reach DNA concentrations that are comparable to DNA concentrations obtained from swabs (see below). Thermocycling used a touchdown approach: one 2-min step at 95°C, 30 cycles of 30 s at 95°C, 30 s at 65°C to 59°C, 90 s at 72°C and one final 5-min step at 72°C before incubation at 15°C. The touchdown protocol implies that the first cycle had a hybridization temperature at 65°C, the following six cycles with hybridization temperatures decreasing down to 59°C by steps of 1°C, and the remaining 24 cycles at 59°C. Amplification success was evaluated by electrophoresis and BET visualization of 5µl of amplification products in 1.75% agarose gels run 30 min at 100V in TAE buffer.

Twelve PCR products issued in balanced numbers (n=4) from muscles, buccal and haemolymph swabs from different individuals were then sent to GenoScreen (https://www.genoscreen.fr/) for Sanger sequencing. Sequencing was performed from both ends.

### Results

—Genomic DNA was detectable in all samples. As expected, it was more concentrated (but this concentration was also more variable) in samples obtained from muscles (mean +/- standard error of the mean: 181,1+/-15,1 ng/µl) as compared to buccal and haemolymph swabs (14.5+/-1.4 ng/µl and 37.1+/-3 ng/µl, respectively). These results also show that there is on average more than double DNA when swabing haemolymph than when swabing the buccal region.

All DNA samples obtained from muscle and haemolymph yielded a visible band at the expected size (202 bp), resulting in a 100% +/- 0% amplification success of this particular locus for these samples. By contrast, amplification with the DNA samples obtained from buccal swabs was less successful (61%, binomial confidence interval: [47%;73%]).

As explained above, we sequenced twelve samples from the successful amplifications. These samples were chosen to represent 12 different individuals and the three types of sampling (muscle, haemolymph and buccal swabs), with four samples each.

The twelve samples yielded high quality sequences. Oficialdegui et al. (2019) found only one haplotype among the 20 individuals they sequenced for a 701 bp fragment while we obtained three different haplotypes from 12 individuals over a 202 bp fragment (data not shown). The first striking result is thus that our pond in Rennes (Outgroup crayfish in the main text) harbours much more genetic diversity than the entire marsh of Brière. The three haplotypes we observed were not in equal frequencies. A first haplotype, obtained from one individual only, cannot be distinguished from Hap_04 and Hap_11 from Oficialdegui et al. (2019). Hap_11 is the only haplotype found in Brière. The two other haplotypes, found in 4 and 7 individuals, belong to the group defined by Hap_09 in Oficialdegui et al. (2019). This haplotype is found in high frequency in Louisiana but also in low frequency in Spain and France.

### Conclusion

—(1) Genomic DNA can reliably be obtained from haemolymph captured on sterile cotton swabs when PIT-tagging crayfish. (2) Crayfish from the pond situated in Rennes display more genetic variability than Brière, all sequences but one tracing an origin that is different from what has been described for the Brière region References

## APPENDIX 3. Trapping devices used in the mesocosms.

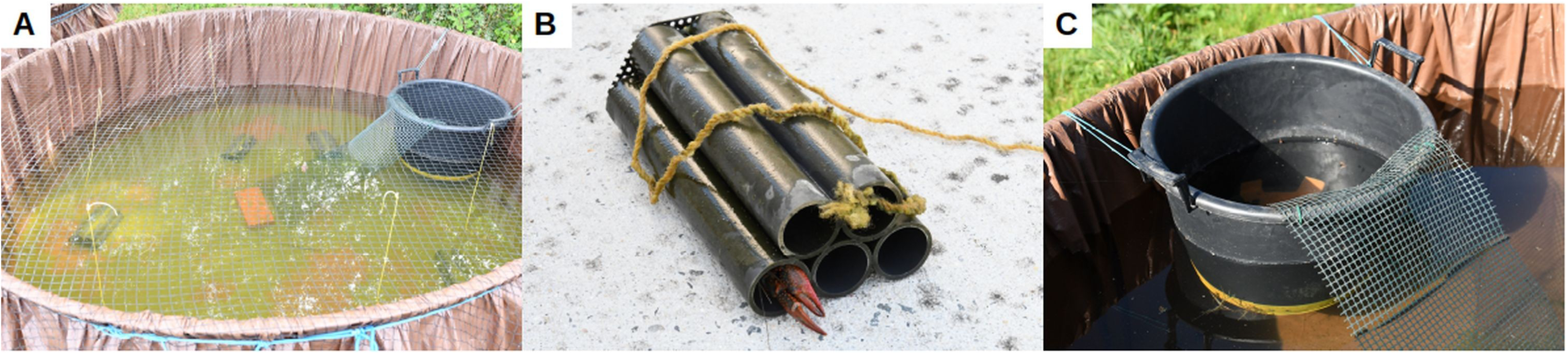

A: Top view of an experimental mesocosm showing artificial refuge traps (ARTs), an exit trap, and algal sediments at the bottom. The brick in the middle of the mesocosm secured the net that served as a ramp to the exit trap. B: Details of an ART with one crayfish inside. C: Details of an exit trap.

## APPENDIX 4. Link between air and water temperature.

Cross-correlation between hourly air temperature at a 5 km-distant weather station and hourly water temperature in six of the 28 the experimental mesocosms. The horizontal blue, dotted lines show the values beyond which the autocorrelation is significantly different from zero at a 5 % risk. Water temperature was highly correlated with air temperature and maximal at a 5-hour lag.

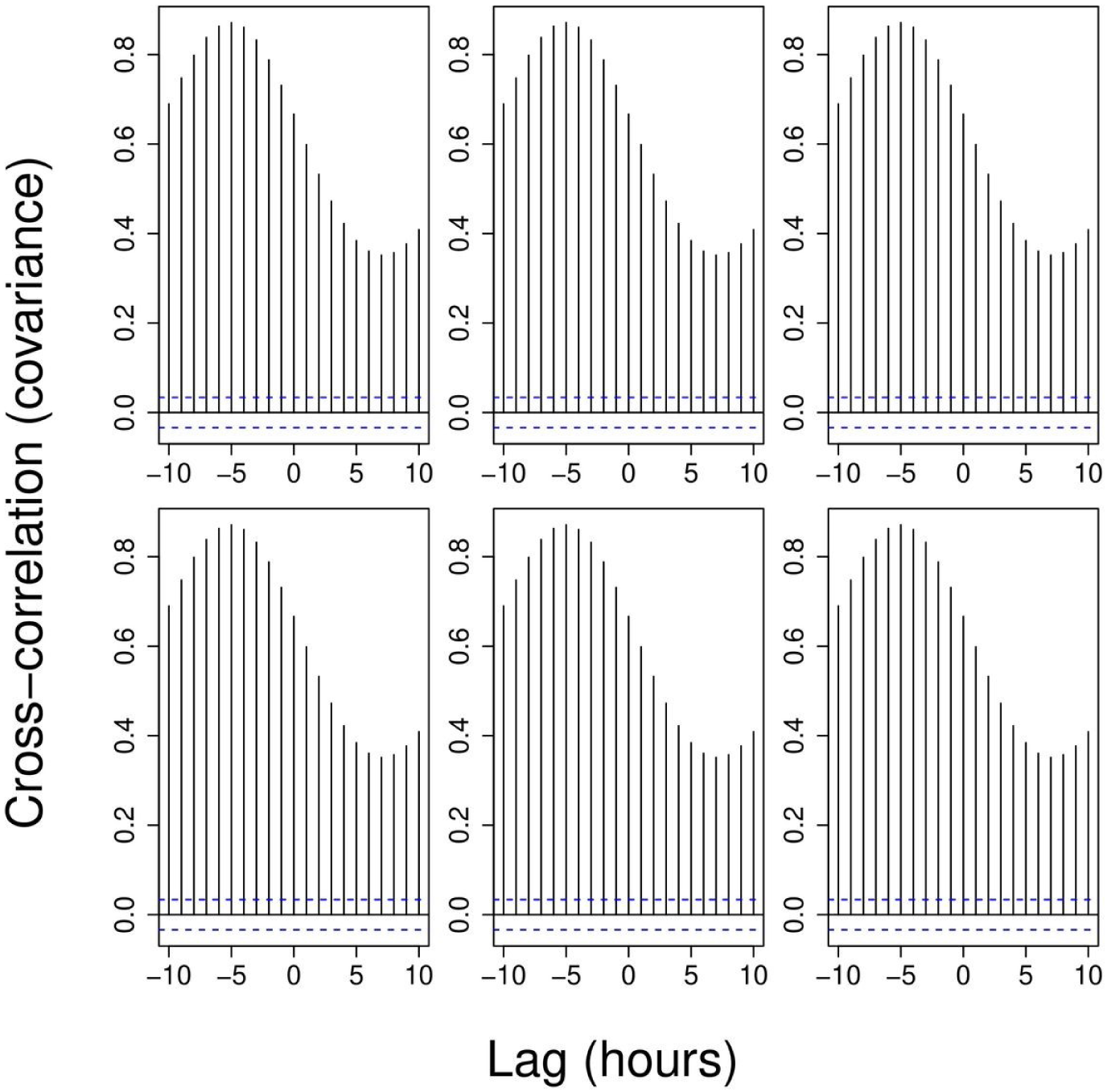

## APPENDIX 5. Parameter estimates from the multistate mark-recapture model using uniform priors on the [- 10,10] interval for regression parameters in Eq. 3.

For more details, see Table 2 legend in the main text.

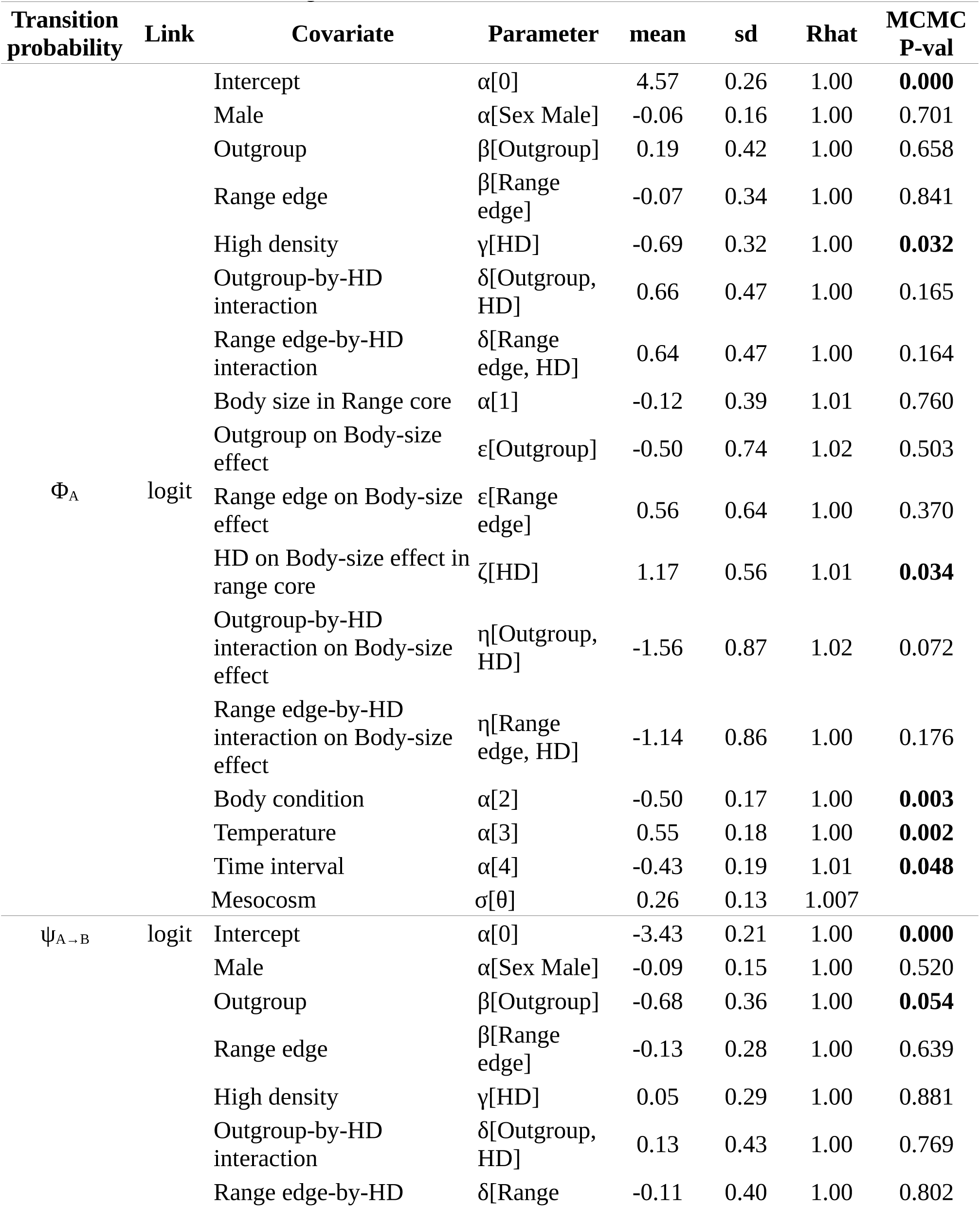
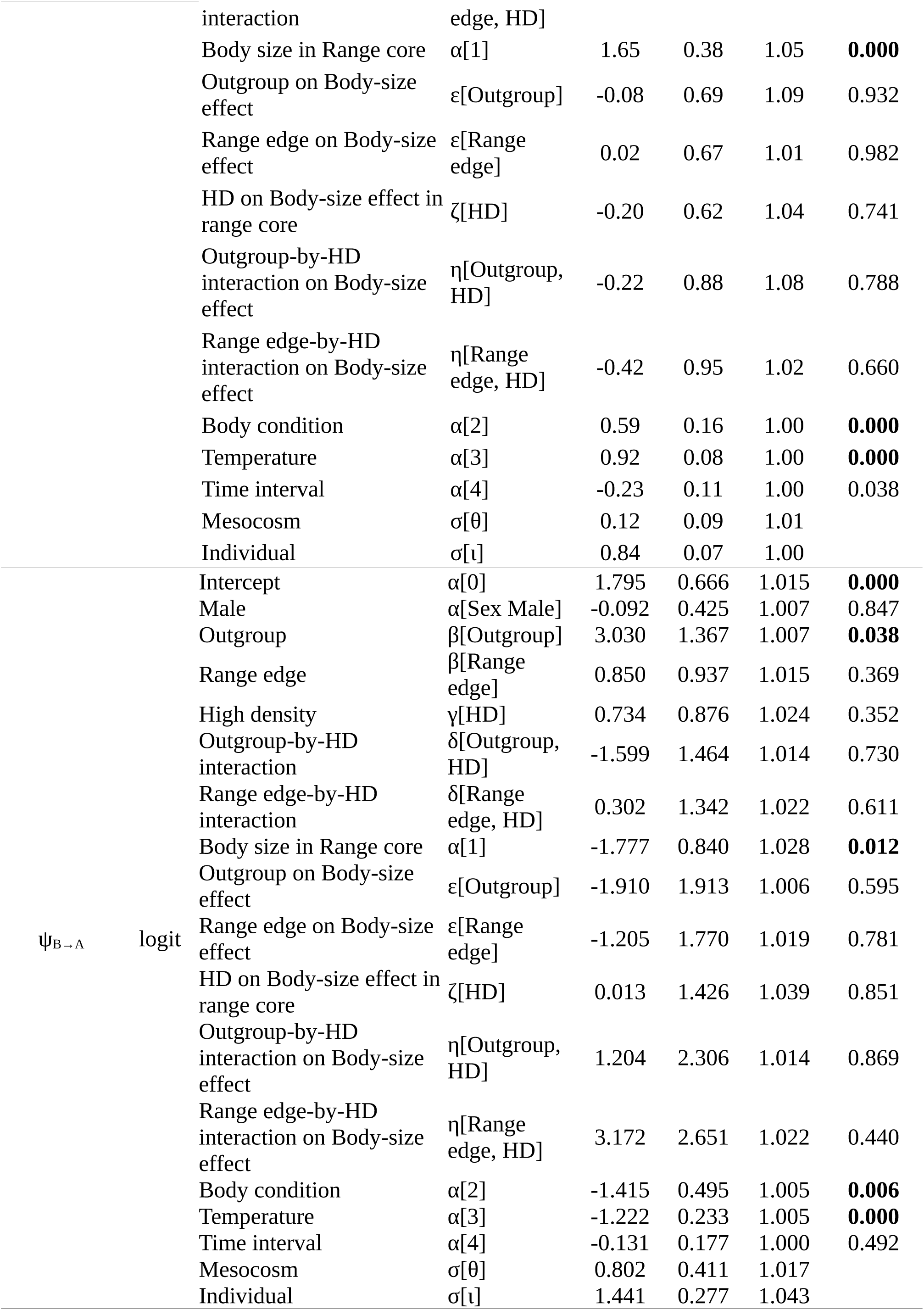

## APPENDIX 6. Densities of prior (yellow bars) and posterior (blue bars) distributions for all main parameters in Eq. (3).

The green colour shows overlap areas. HD = high-density treatment.

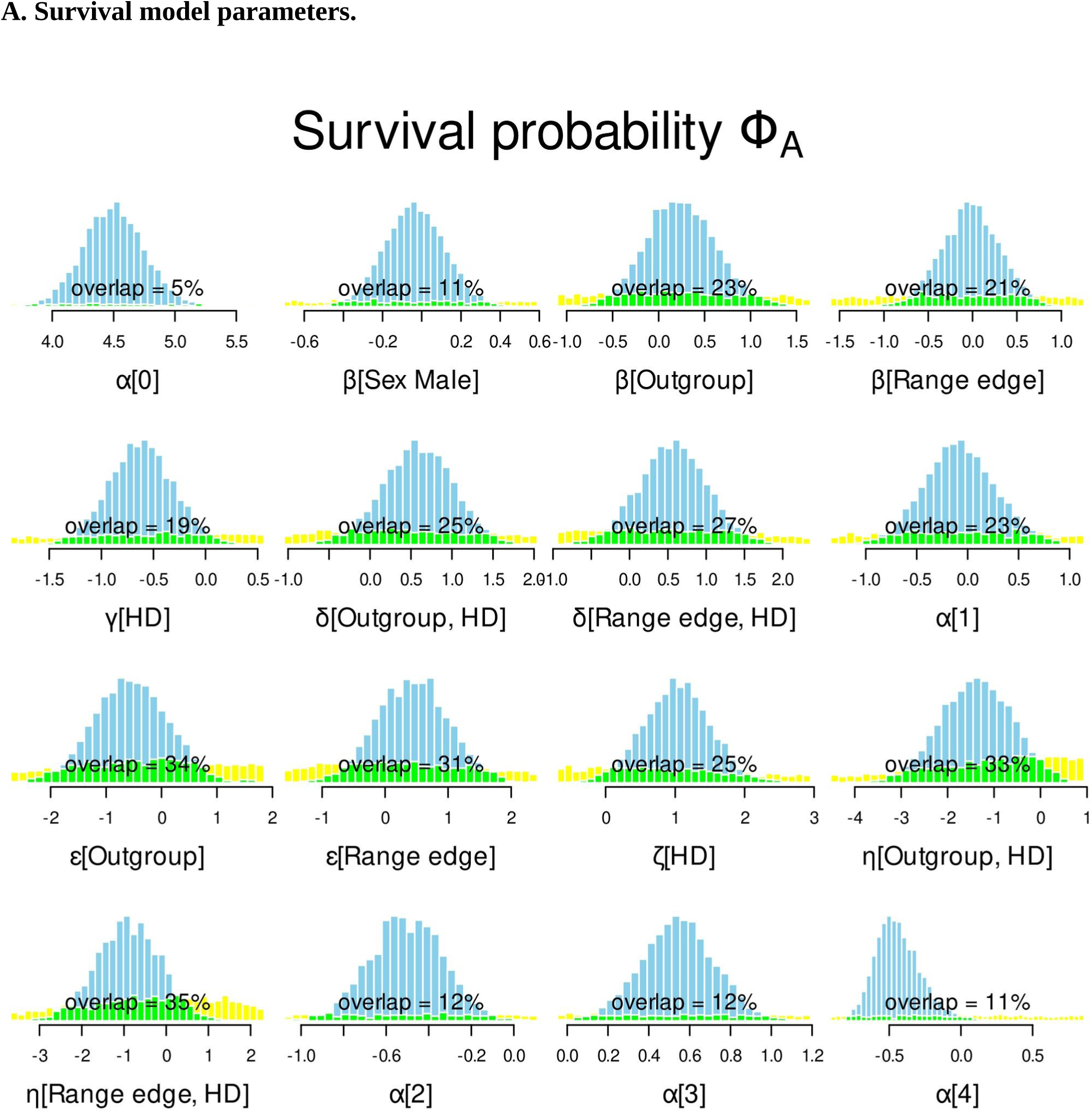
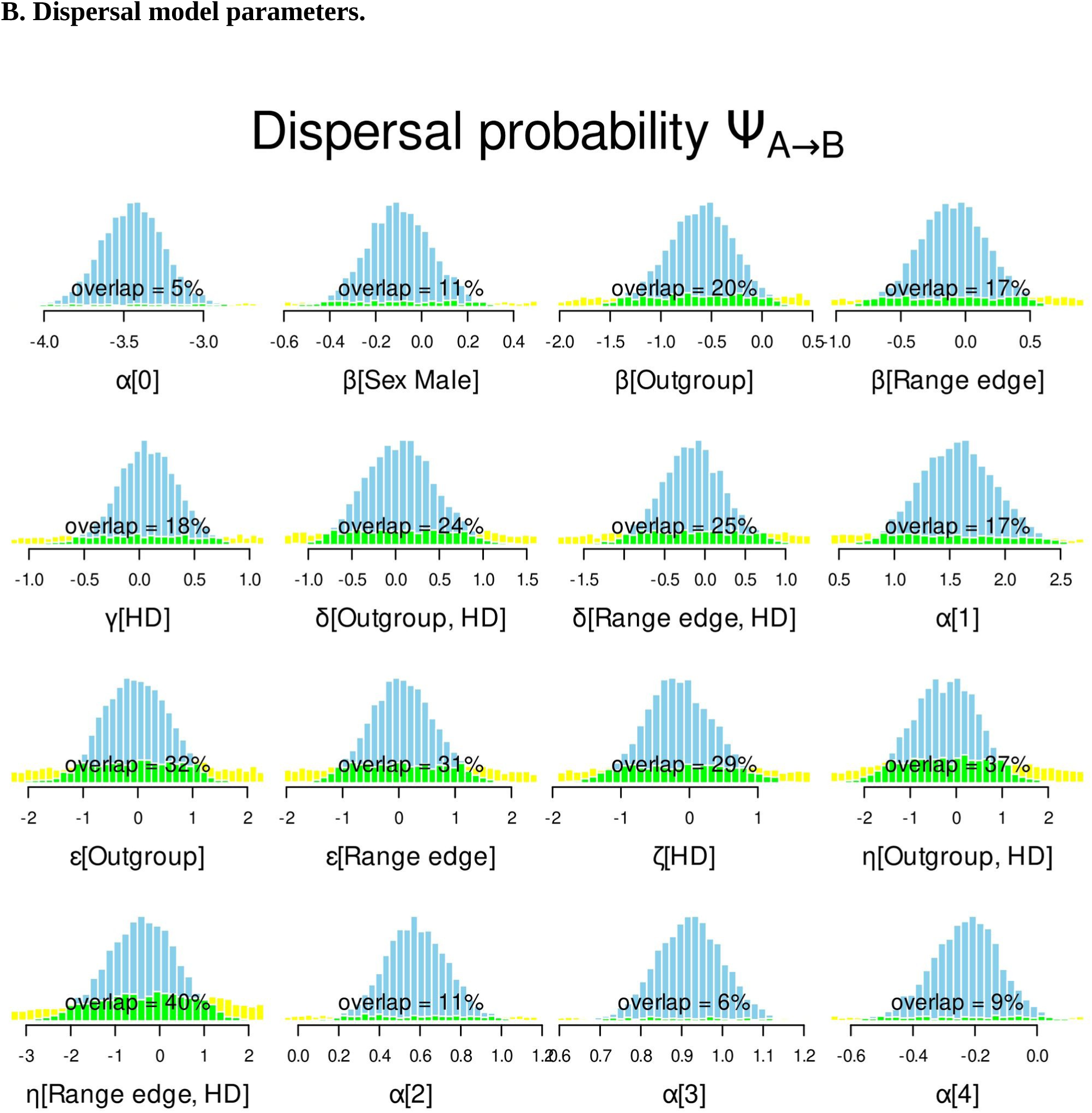
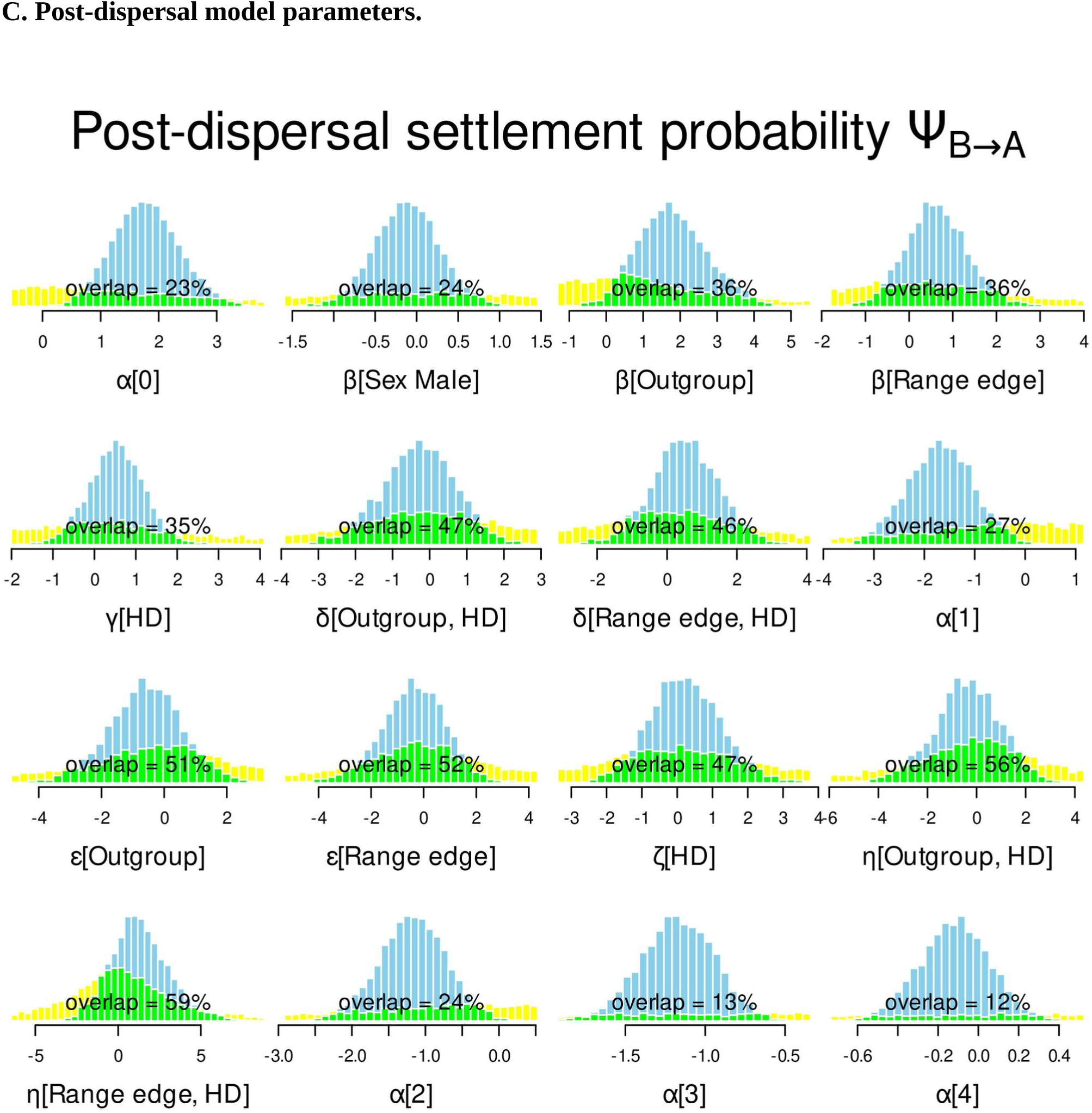

## APPENDIX 7. Parameter estimates for a model including only main effects on crayfish probabilities of survival Φ_A_, dispersal ψ_A→B_ and post-dispersal settlement ψ_B→A_.

For more details, see Table 2 legend in the main text.

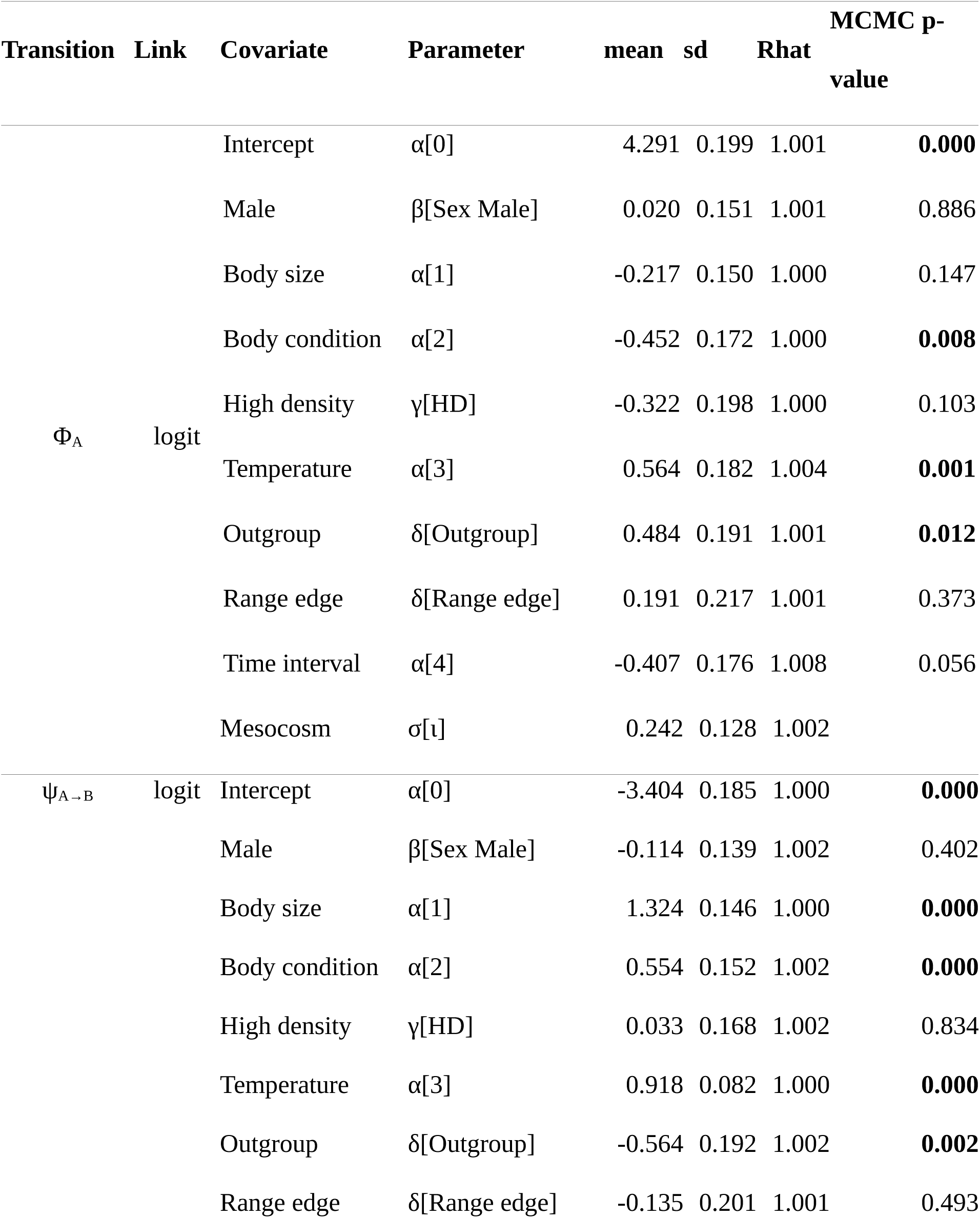
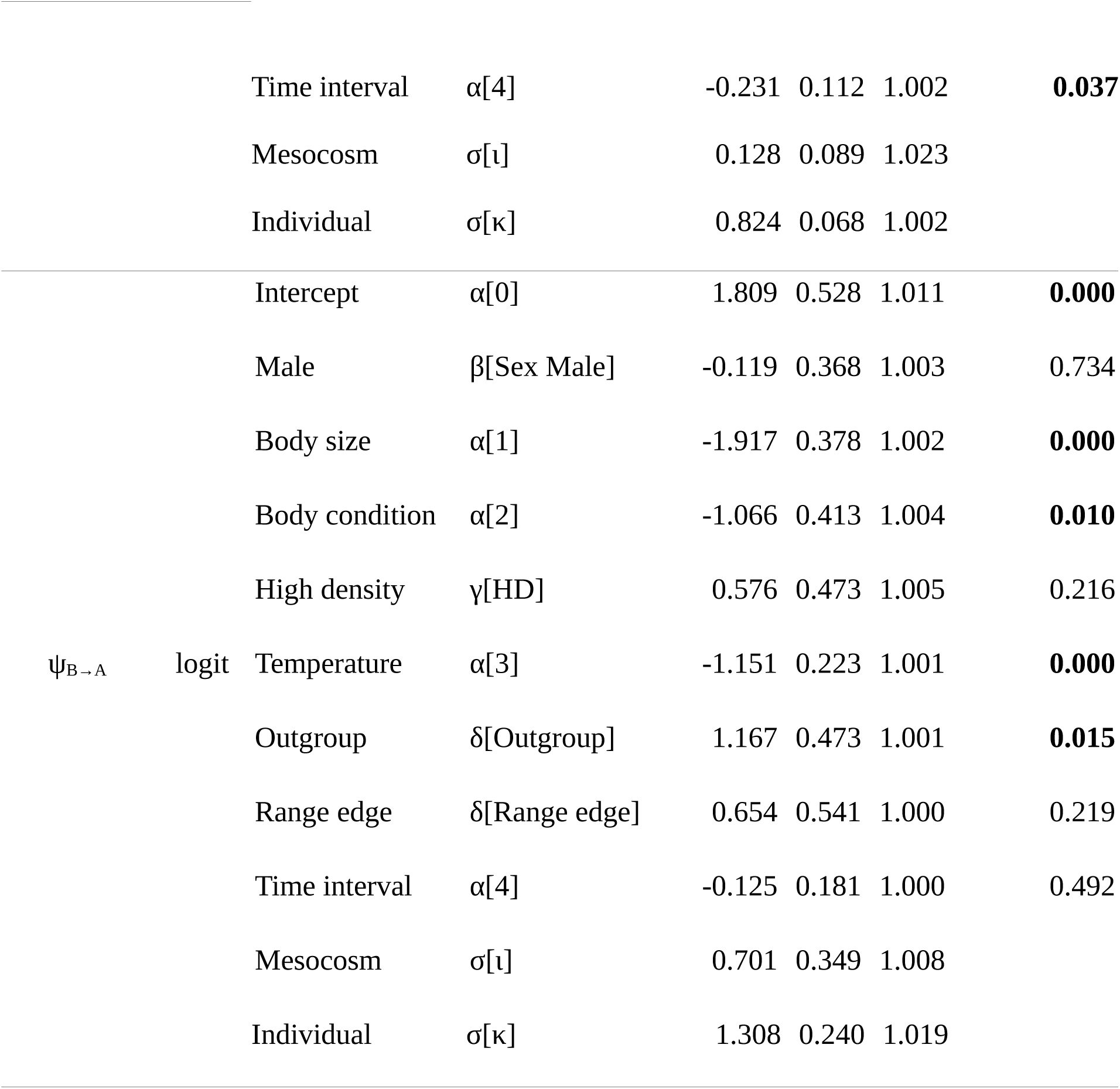

## APPENDIX 8. Effects of stocking density and experiment duration on crayfish survival rate in aquaculture studies.

Density varied from 1 to 1200 individuals m^-2^, and duration from 3 to 306 days. N = 436 observations from 29 studies. Species included: *Astacus leptodactylus:* n = 107, *Austropotamobius pallipes:* n = 32, *Cherax destructor:* n = 55, *Cherax quadricarinatus:* n = 110, *Pacifastacus leniusculus:* n = 45, *Pontastacus leptodactylus:* n = 3, *Procambarus clarkii:* n = 84. Relationship modelled from a beta GLMM (logit link) with log(density) and duration and their interaction as fixed effects, and with the study as random intercept in the glmmTMB library of R (Brooks et al., 2017). The effect of crayfish species increased model AIC and was thus discarded. Reference list cited below.

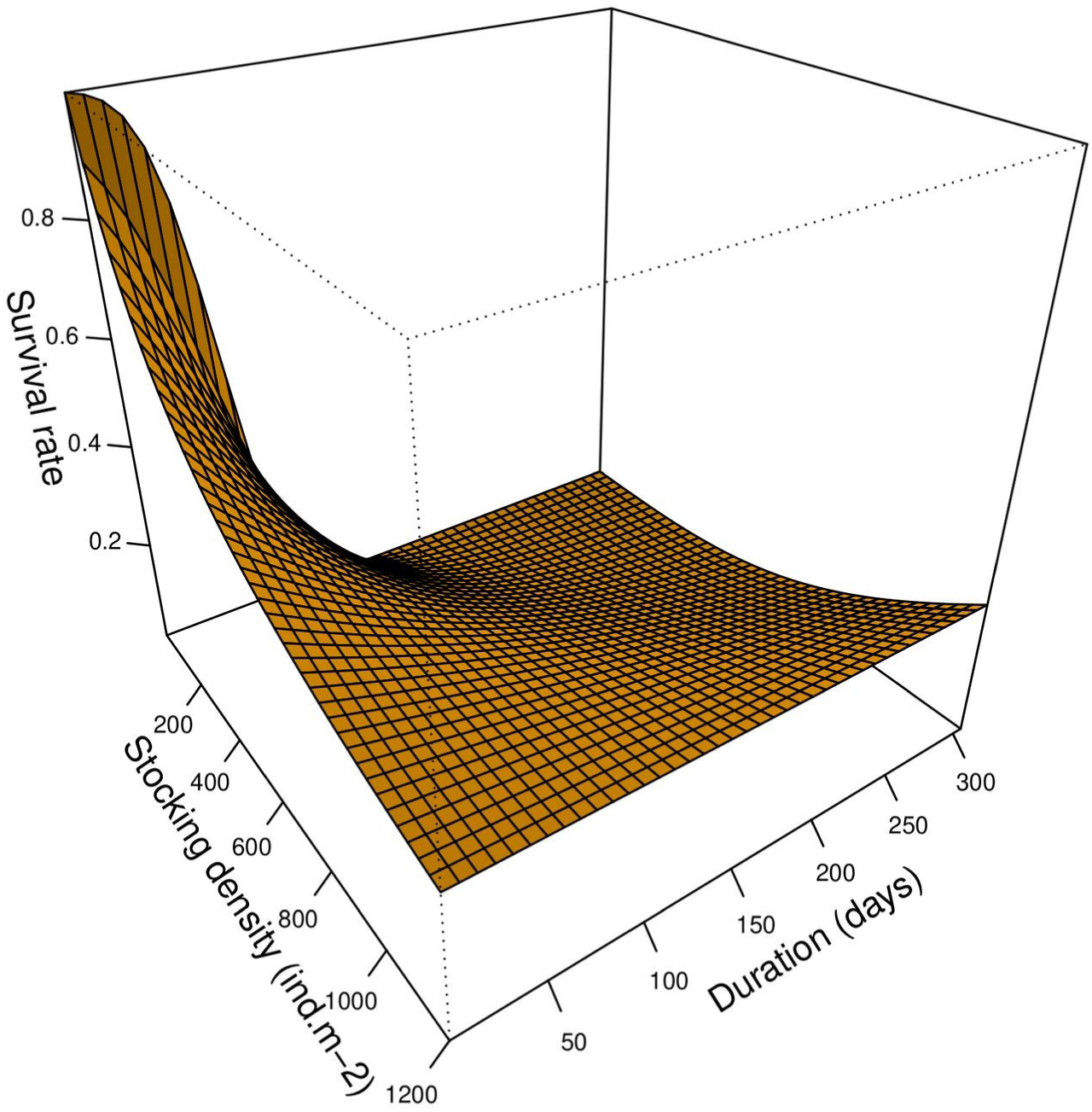

## APPENDIX 9. Predicted crayfish survival and overland dispersal probabilities by 2071-2100 under the SSP1-2.6 and SSP5-8.5 scenarios of IPCC.

Predictions were formed from parameter estimates for the multistate mark-recapture model (Table 2), for an average-sized Range-core female at low crayfish density. Annual mean reference temperature was 10.7°C (“Historical” situation), which is average summer (May-September) temperature in Rennes City France for the 1981-2010 period (https://fr.wikipedia.org/wiki/Rennes).

|  | Historical | SSP1-2.6 scenario |  | SSP5-8.5 scenario |  |
| --- | --- | --- | --- | --- | --- |
|  |  | Low | High | Low | High |
| Monthly survival probability | 0.84 | 0.86 | 0.87 | 0.87 | 0.89 |
| Time to dispersal probability = 1 (days) | 111 | 95 | 81 | 83 | 62 |

## Footnotes

1 https://fr.wikipedia.org/wiki/Rennes

